# Improving Hi-C contact matrices using genome graphs

**DOI:** 10.1101/2023.11.08.566275

**Authors:** Yihang Shen, Lingge Yu, Yutong Qiu, Tianyu Zhang, Carl Kingsford

## Abstract

Three-dimensional chromosome structure plays an important role in fundamental genomic functions. Hi-C, a high-throughput, sequencing-based technique, has drastically expanded our comprehension of 3D chromosome structures. The first step of Hi-C analysis pipeline involves mapping sequencing reads from Hi-C to linear reference genomes. However, the linear reference genome does not incorporate genetic variation information, which can lead to incorrect read alignments, especially when analyzing samples with substantial genomic differences from the reference such as cancer samples. Using genome graphs as the reference facilitates more accurate mapping of reads, however, new algorithms are required for inferring linear genomes from Hi-C reads mapped on genome graphs and constructing corresponding Hi-C contact matrices, which is a prerequisite for the subsequent steps of the Hi-C analysis such as identifying topologically associated domains and calling chromatin loops. We introduce the problem of genome sequence inference from Hi-C data mediated by genome graphs. We formalize this problem, show the hardness of solving this problem, and introduce a novel heuristic algorithm specifically tailored to this problem. We provide a theoretical analysis to evaluate the efficacy of our algorithm. Finally, our empirical experiments indicate that the linear genomes inferred from our method lead to the creation of improved Hi-C contact matrices. These enhanced matrices show a reduction in erroneous patterns caused by structural variations and are more effective in accurately capturing the structures of topologically associated domains.

## 1 Introduction

The spatial arrangement of chromosomes plays an important role in many crucial cellular processes including gene transcription [Fraser and Bickmore, 2007, Rennie et al., 2018], epigenetic modification [Grewal and Moazed, 2003], and replication timing [Pope et al., 2014]. This complex structure can be discovered through Hi-C [Lieberman-Aiden et al., 2009], a high-throughput variant of the chromosome conformation capture technique [Dekker et al., 2002], which has become a prevalent tool in the study of genomic organization. The Hi-C process yields read pairs representing spatial contacts between two genomic loci. These contacts can be identified by aligning each end of a read pair to the reference genome. These aligned read pairs facilitate subsequent analyses, such as identifying topologically associated domains (TADs) [Nora et al., 2012, Dixon et al., 2012, De Laat and Duboule, 2013, Filippova et al., 2014, Li et al., 2018], which are contiguous regions on chromosomes with more frequent contacts, and calling chromatin loops [Roayaei Ardakany et al., 2020], which are pairs of genomic loci that lie far apart along the linear genome but are in close spatial proximity.

Hi-C pipelines use the linear reference genome such as Genome Reference Consortium Human Build 38 (GRCh38) as the template against which to align reads. However, these linear references do not incorporate the genetic diversity within populations. Consequently, aligning reads from genomes that diverge from the linear reference genome can result in reads either not aligning at all or being mapped to incorrect genomic locations. This issue is exacerbated when analyzing Hi-C reads from cancer samples, which frequently exhibit structural variations, including copy number variations and substantial translocations. The misalignments, arising due to structural variations, can confound the interpretation of Hi-C data, potentially producing features that may be mistaken for other biological signals, such as chromatin loops [Wang et al., 2020]. Given that read alignment is always the initial step in Hi-C analysis, errors at this stage can proliferate, leading to inaccuracies throughout the downstream analyses. To rectify the Hi-C analysis of cancer cell lines, substantial efforts have been made to develop algorithms to identify structural variations and rearrange the cancer genomes from Hi-C data, sometimes with the help of other data types such as whole genome sequencing to enhance accuracy and precision [Wang et al., 2020, Schöpflin et al., 2022, Khalil et al., 2020, Wang et al., 2021] of Hi-C analysis of cancer cell lines. However, these steps still rely on the linear reference genome.

The concept of pan-genome has been introduced to address the shortcomings of linear references. The pan-genome is a collection of DNA sequences that incorporates both common DNA regions and sequences unique to each individual [Gong et al., 2023, Wang et al., 2022]. These DNA sequences can be represented by genome graphs, which are graph-based data structures amalgamating the linear reference alongside genetic variations and polymorphic haplotypes [Ameur, 2019]. Numerous computational techniques have been published in the domain of genome graphs, addressing various aspects such as efficient genome graph construction [Garrison et al., 2018, Qiu and Kingsford, 2021, Hickey et al., 2023, Garrison et al., 2023, Pandey et al., 2021], graph-based genome alignment [Rakocevic et al., 2019, Kim et al., 2019, Rautiainen and Marschall, 2020, Sirén et al., 2021] and graph-based structural variation and haplotype analyses [Chin et al., 2023, Hadi et al., 2020, Eggertsson et al., 2019, Chen et al., 2019]. These studies have substantiated that genome graphs can enhance the analysis of genome sequencing data. Moreover, Liao et al. [2023] has illustrated the potential of genome graphs in improving the analysis of various other data types, including RNA-seq, ChIP-seq, and ATAC-seq. However, there has been no research exploring the enhancement of Hi-C analyses through the use of genome graphs.

Leveraging genome graphs as the reference can enhance the accuracy of mapping Hi-C reads. However, a challenge arises from the noisy information in the graphs post-read alignment, attributed to structural variations present in the genome graphs but absent in the actual linear genome of the Hi-C sample. Additionally, these graphs are not immediately applicable for subsequent standard Hi-C analysis steps like TAD identification [Filippova et al., 2014, Wang et al., 2017, Li et al., 2018] and chromatin loop calling [Ay et al., 2014], given their inherent dependence on linear genomes and the corresponding Hi-C contact matrices, the two-dimensional matrices representing chromosomal interactions.

A critical component to address this is to infer the appropriate, sample-specific linear genome from Hi-C reads mapped on genome graphs. These inferred genomes, which are more congruent with the Hi-C samples’ unknown ground truth genomes than traditional linear reference genomes, account for polymorphisms and structural variations specific to the given sample. By using these reconstructed genomes to create Hi-C contact matrices and incorporating these matrices into subsequent analyses, we can enhance the precision of Hi-C studies. This approach offers a more customized and sample-specific genomic representation, addressing the shortcomings inherent in using standard linear reference genomes.

In this work, we investigate the problem of genome sequence inference from Hi-C data on directed acyclic genome graphs. We propose a novel problem objective to formalize the inference problem. To infer the genome, we choose the best source-to-sink path in the directed acyclic graph that optimizes the confidence of TAD inference on the genomes. We show that optimizing this objective is NP-complete, a complexity that persists even with directed acyclic graphs. We develop a greedy heuristic for the problem and theoretically show that, under a set of relaxed assumptions, the heuristic finds the optimal path with a high probability. To ensure practical applicability to real Hi-C datasets, we also develop the first complete graph-based Hi-C processing pipeline. We test our processing pipeline and genome inference algorithms on cancer Hi-C samples K-562 and KBM-7. Results on these samples show that compared to using the traditional linear reference genomes, the linear genomes inferred from our method create improved Hi-C contact matrices. These enhanced matrices exhibit fewer errors caused by structural variations and are more effective in accurately capturing the structures of TADs, attesting to the algorithm’s potential to enhance the precision and reliability of genomic studies. The source code is available at https://github.com/Kingsford-Group/graphhic.

## 2 Inferring the sample genome from Hi-C data with genome graphs

### 2.1 Preprocessing of Hi-C reads via a novel graph-based Hi-C pipeline

Typical Hi-C processing pipelines, such as HiC-Pro [Servant et al., 2015], mainly consist of two steps: (i) aligning each end of raw read pairs to the linear reference genome; (ii) constructing a two-dimensional contact matrix that describes the interactions between pairs of genomic regions. A contact matrix is derived from the alignment results, wherein each entry contains the number of read pairs between two genomic bins — intervals with a fixed length such as 10 kilobases. This contact matrix is used as an input for downstream analyses, such as TAD identification and loop calling. However, current Hi-C analysis pipelines are unable to process data when genome graphs are used as the reference instead of linear reference.

We develop a novel graph-based Hi-C processing pipeline shown in Figure 1. Our Hi-C pipeline is composed of four steps. First, it constructs directed acyclic genome graphs either from various DNA sequences or from the linear reference genome coupled with population variants represented in Variant Call Format (`.vcf`) files. Each node in the genome graph represents a DNA subsequence. Second, the pipeline performs graph-based alignment to map Hi-C reads onto the paths of the genome graphs. In the third step, we prune and contract the resulting genome graph by removing paths with no reads mapped and contracting nodes into larger genomic bins, which greatly improves the computational efficiency of the following steps. Subsequently, the pipeline builds a contact matrix *M* based on the pruned genome graph. Each dimension of the matrix represents nodes of the pruned graphs, and each matrix entry is the number of read pairs with ends mapped to the corresponding nodes. The nodes are ordered in their topological order in the genome graph. An exhaustive delineation of each stage of our pipeline is provided in Appendix A.

**Figure 1.**
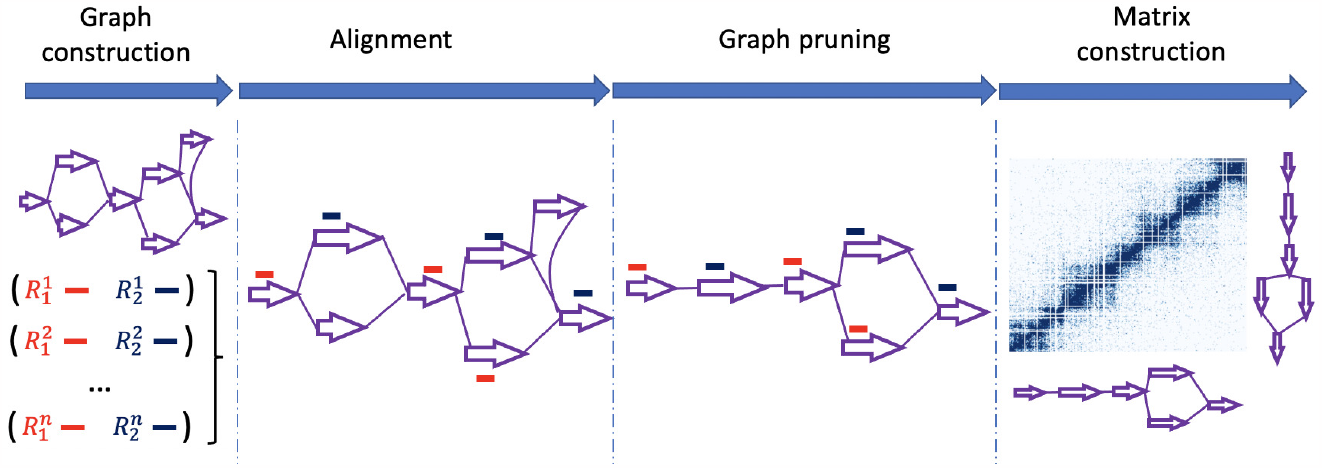
The workflow of our graph-based Hi-C processing pipeline. Each red line represents one end of a read pair, and each blue line represents the other end.

### 2.2 Problem definition of genome inference

Given a directed acyclic genome graph *G* with a source node *s* and a sink node *t*, and the contact matrix *M* derived from the graph-based Hi-C pipeline (Section 2.1), the objective of genome inference is to find a *s*-*t* path in *G* such that the concatenated DNA sequences represented by nodes in the selected path is the most similar sequence to the actual genome of the Hi-C sample. Ideally, two primary criteria should be fulfilled by this reconstructed path: (i) it should encapsulate as many mapped Hi-C read pairs as feasible, and (ii) the distribution of these mapped pairs ought to echo the distinctive spatial structures of chromosomes, especially the topologically associated domains (TADs or “domains” for brevity). Motivated by these prerequisites, our approach toward genome inference encompasses a simultaneous inference of the *s*-*t* path and the corresponding TADs from *G*.

Let *P*_*st*_ be the collection of all *s*-*t* paths in *G*, and let *D*_*p*_ be a set of domains along path *p* ∈ *P*_*st*_. The *i*-th domain on path *p* is defined as a subpath 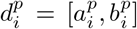 that starts at node 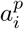 and ends at 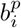, where 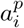 and 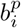 are nodes on *p*. We require that domains of one path do not overlap with each other. Furthermore, the boundaries of two consecutive domains 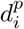 and 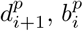and 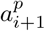, are two nodes connected with an edge on path *p*. The first and the last domain are 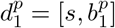 and 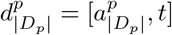.

We infer the *s*-*t* path representing the actual genome from a Hi-C sample by solving the following problem:

#### Problem 1.

We are given a directed, acyclic graph *G* = (*V, E*) with a source node *s* and a sink node *t*, a pre-computed function *μ* : ℕ → ℝ_≥0_, a float value *γ* ≥ 0, and a cost function *c* : *V* × *V* → ℝ_≥0_ that maps every pair of nodes to a non-negative cost. *c* is symmetric in a sense that *c*(*v, v*^′^) = *c*(*v*^′^;, *v*). The goal is to find a *s*-*t* path *p* = {*v*_1_, *v*_2_, …, *v*_|*p*|_ } over all *s*-*t* paths and a set of consecutive domains *D*_*p*_ on *p* that optimize

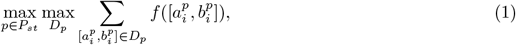

where *f* is defined as:

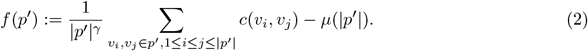

The cost function *c*(*v*_*i*_, *v*_*j*_) can be described by entries in the contact matrix *M* (*v*_*i*_, *v*_*j*_) for node *v*_*i*_ and node *v*_*j*_. *f* quantifies the quality of a TAD as the normalized number of interactions within the subpath *p*^′^. 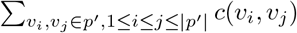 in equation (2) is the total number of interactions between nodes that are both in path *p*^′^.

The total number of interactions is normalized by two factors. First, the total number is zero-centered by a pre-computed *μ*(|*p*^′^|), which is the expected interaction frequency within a path with |*p*^′^| nodes. Then, it is normalized by he number of nodes in *p*^′^ (|*p*^′^|) scaled by a factor of *γ*. This normalization prevents the identification of TAD domains with excessively large sizes. Larger values of *γ* typically lead to finding smaller domains.

### 2.3 The hardness of the problem

The objective function (1) of Problem 1 is derived from that of Filippova et al. [2014]. However, Filippova et al. [2014] employs polynomial-time dynamic programming to infer TADs based on reads mapped to a linear reference, while our work requires the concurrent inference of both the TAD domains (denoted as *D*_*p*_) and the sample’s linear genome (represented by the path *p*) directly from *G*. We show that this increased complexity results in NP-completeness of Problem 1.

#### Theorem 1.

Problem 1 is NP-complete.

We prove Theorem 1 by reducing from the Path Avoiding Forbidden Pairs (PAFP) problem, which has been confirmed to be NP-complete even in directed acyclic graphs [Gabow et al., 1976, Kolman and Pangrác, 2009].

#### Problem 2

(Path avoiding forbidden pairs problem [Kolman and Pangrác, 2009]). Given a graph G = (*V, E*) with two fixed vertices *s, t* ∈ *V* and a set of pairs of vertices *F* ⊂ *V* × *V*, find a path from *s* to *t* that contains at most one vertex from each pair in *F*, or recognize that such path does not exist.

The core concept of the proof involves transforming a graph instance from PAFP into a novel graph instance of Problem 1. This transformation is done so that the objective (1) attains a specific value in the new instance if and only if a path avoiding all forbidden pairs exists in the original graph instance. The details of the proof are in Appendix B. This hardness result motivates the development of practical heuristics for the problem.

### 2.4 Computation of the *μ* function

Filippova et al. [2014] demonstrated a method for efficiently pre-computing *μ* on the linear reference genome. Nevertheless, as we discuss in Appendix F, calculating *μ* in the context of genome graphs poses a more complex challenge. Consequently, we propose a different strategy to estimate *μ*. Generally, samples from normal cell types bear a greater resemblance to the linear reference genome compared to those from cancer samples. Hence, we select Hi-C data from a normal sample, process it using the linear reference genome, and calculate its *μ* function using the same approach as Filippova et al. [2014]. This function is denoted as *μ*_0_. It is evident that the sequencing depth of the Hi-C data can influence the value of the *μ* function. Therefore, when analyzing new Hi-C data, we estimate its *μ* function as follows:

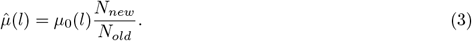

Here, *N*_*old*_ refers to the total count of Hi-C read pairs from the original normal sample, while *N*_*new*_ indicates the total count of Hi-C read pairs from new Hi-C data.

### 2.5 Graph-based dynamic programming algorithm

We use a dynamic program, computed in the topological ordering of the nodes, to solve the Problem 1:

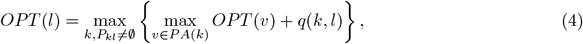

where *OPT* (*l*) is the optimal solution for objective (1), applied to the subgraph induced by node *l* along with all nodes with a topological order less than that of *l* within *G. P*_*kl*_ denotes the collection of all paths from node *k* to node *l* in *G*, and *PA*(*k*) is the set of parent nodes of *k*. max_*v*∈*P A*(*k*)_ *OPT* (*v*) = 0 if *k* has no parent node. *q* is a function that maximizes over all paths between *k* and *l*:

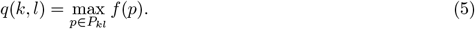

We use a standard backtracking strategy, shown in Algorithm 3 in Appendix C, to reconstruct the optimal path *p*_*opt*_ from the dynamic program. The reconstructed path *p*_*opt*_ is the inferred genome we want.

We prove that *OPT* (*t*) is indeed the optimal solution for Problem 1.

#### Proposition 1.

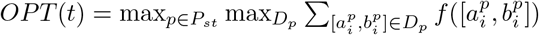

The proof is in Appendix D. However, this does not provide a polynomial time algorithm. As we show in Appendix E, computing the function *q* in (5) is NP-complete. Moreover, the NP-completeness of computing *q* is not attributable to the exclusive definition of function *f* as outlined in (2). We show in Appendix E that computing *q* remains NP-complete under various definitions of *f* . Therefore, it is hard to solve the dynamic programming objective shown in (4), which is consistent with the hardness conclusions in Section 2.3. This provides a focus for developing heuristics.

### 2.6 Heuristics for computing *q*

We propose a novel heuristic algorithm, detailed in Algorithm 1, to compute the function *q*(*k, l*) for any node pair (*k, l*). The central principle behind this algorithm is that a node situated between nodes *k* and *l* (in topological order) that has more interactions with other nodes is more likely to be a part of the path connecting *k* to *l* that maximizes the function *f* . Consequently, we sort the nodes in descending order based on their cumulative interactions with other nodes (line 6 of Algorithm 1) and progressively add nodes from the highest to the lowest interactions until a *k*-*l* path is established.

#### Algorithm 1 *q*(*k, l*) computation

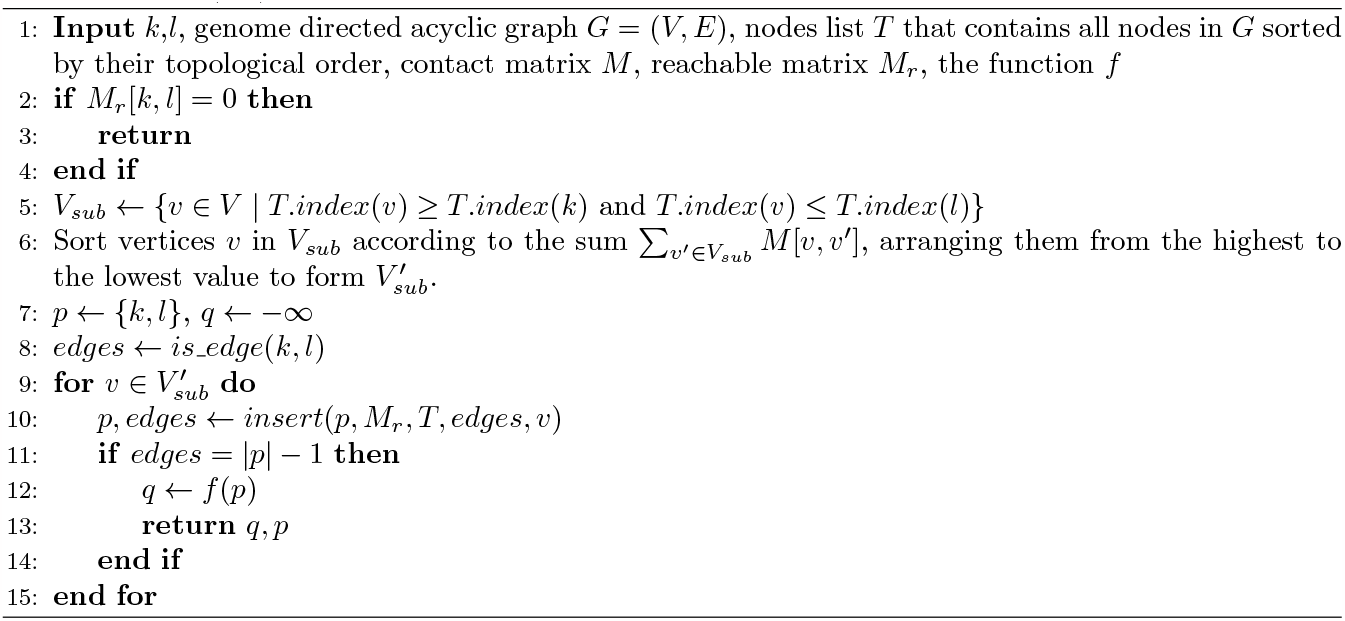

Specifically, we employ the following functions and data structures within Algorithm 1 to enhance the algorithm’s efficiency:

- reachable matrix *M*_*r*_, where *M*_*r*_[*i, j*] = 1 if there exists a path from node *i* to node *j* in the directed acyclic genome graph *G*, otherwise *M*_*r*_[*i, j*] = 0.
- *is edge*(*k, l*), which returns 1 if there is an edge from *k* to *l* in *G*, otherwise it returns 0.
- *insert*(*p, M*_*r*_, *T, edges, v*), of which the pseudo-code is provided in Algorithm 4 in the appendix. This function contains the following steps:
  - Given a node set *p* which encompasses all nodes already incorporated and are topologically sorted, the function determines whether there exists a path in *G* that includes all nodes in *p* ∪ {*v*}. Such a path may include additional nodes that are not in *p* ∪ {*v*}. This step can be efficiently achieved with the help of *M*_*r*_ and a balanced tree structure such as AVL tree [Foster, 1973], of which the details are introduced in Appendix G.2.
  - If the aforementioned path exists, the node *v* is then inserted into *p* according to the topological ordering (function *update* in line 7 of Algorithm 4).
  - The function also updates an integer variable *edges* (line 8 of Algorithm 4), which keeps track of how many neighboring nodes in *p* have edges in *G*.

The *insert* function yields a revised node set *p* and an updated value for *edges* (line 10 of Algorithm 1). A legitimate path in graph *G* is formed by the nodes in *p* if and only if the condition *edges* = |*p*| − 1 is met (line 11 of Algorithm 1). Once a path is established, we compute the function value *f* (*p*) and use it as the value of *q* (line 12 of Algorithm 1).

We have the following result on the time complexity of our heuristic algorithm:

#### Theorem 2.

The total time complexity for Algorithm 1 and the dynamic program using the heuristic Algorithm 1 are respectively 𝒪 (|*V* |^2^) and 𝒪 (|*V* |^4^), where |*V* | is the number of nodes in the graph.

The proof is in Appendix G.2. In practice, the time complexity is still too high for long chromosomes. To address this, as detailed in Section 2.8, we implement additional practical strategies to further decrease the algorithm’s time complexity.

### 2.7 Accuracy of the heuristic algorithm

Let 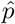 represent the path from node *k* to node *l* as predicted by Algorithm 1, and let *p*_*gt*_ denote the “ground truth” path, defined as 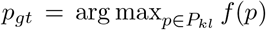. In an ideal scenario, a heuristic algorithm would ensure that, for any specified DAG *G* and any interaction distribution present on *G*, the value 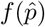 closely approximates *f* (*p*_*gt*_). However, as we demonstrate in Appendix G.3, it is possible to create an example where the discrepancy between 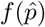 and *f* (*p*_*gt*_) can be infinitely large, indicating that our heuristic algorithm does not offer a bounded approximation in the worst-case scenario.

However, within the scope of Hi-C analysis, the distribution of interactions on a genome graph is not arbitrary. Conceptually, each interaction, represented by a pair of nodes, stems from two primary sources: (a) the “ground truth” source. Both nodes of the interactions from this source lie on the ground truth path *p*_*gt*_. Interactions from this source are informative when constructing 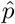. (b) the “noise” source, which accounts for interactions arising due to various systematic biases such as sequencing errors, mapping errors, etc. In this scenario, the interactions are not necessarily confined to the path *p*_*gt*_. Under mild assumptions, we propose a theoretical framework that more accurately reflects the real-world Hi-C situation, and demonstrate that with high probability the output path 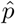 is equivalent to *p*_*gt*_, as long as the number of mapped read pairs is at least Ω(|*p*_*gt*_| log |*V* |). Although the number of total nodes |*V* | in the graph can be large, the required number of read pairs for a successful inference is only proportional to the logarithm of it. The details of this theoretical analysis can be found in Appendix G.4. We observe that in practice, this criterion regarding the number of read pairs is readily met. For instance, in our experiments, the graph has approximately 5 × 10^5^ nodes, and the total number of mapped read pairs is around 3 × 10^8^. This result provides some theoretical justification for the choice of the heuristic in Algorithm 1.

### 2.8 Practical improvements to efficiency and accuracy

In practice, we introduce two modifications to our heuristic algorithm to enhance its accuracy and speed. First, given that the size of TADs typically does not exceed 3Mb [Bonev and Cavalli, 2016], we implement an additional heuristic adjustment to the dynamic program. When calculating the function *q*, we restrict our consideration to paths where the combined length of the DNA sequences on the nodes is under 3Mb. This heuristic modification means that the dynamic program represented by equation (4) would be transformed as follows:

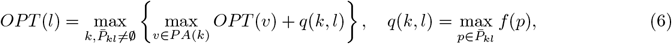

where 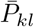 is the collection of paths from *k* to *l* that satisfy the constraint described above. Let *L* denote the largest length of the paths in 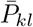, where length here refers to the number of nodes; generally, 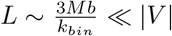|. Now the time complexity of the dynamic program when using the heuristic algorithm for *q* becomes 𝒪 (|*E*||*V* | + *L*^4^), where 𝒪 (|*E*||*V* |) comes from computing the reachable matrix *M*_*r*_.

Second, our empirical observations suggest that for most node pairs (*k, l*), computing *q* using Algorithm 1 is quite effective. Nonetheless, this method might not adequately capture the signals of large deletions. To mitigate this, we have refined Algorithm 1, as detailed in Algorithm 5 in Appendix G.5. In this adjustment, for each node pair (*k, l*), we initially execute a node-weighted shortest path algorithm (where each node’s weight is determined by the length of its corresponding DNA sequence) to identify a path *p*_*base*_ and calculate its score (lines 5-6 of Algorithm 5). Subsequently, Algorithm 1 is applied; if the path *p* derived from Algorithm 1 surpasses the score of *p*_*base*_, *p* is returned, otherwise *p*_*base*_ is the selected path.

The shortest path algorithm for directed graphs with nonnegative weights has a time complexity of 𝒪 (|*E*| + |*V* | log(|*V* |)). Consequently, the overall time complexity for Algorithm 5 to estimate *q* remains 𝒪 (|*V* |^2^) (or 𝒪 (|*L*|^2^) if we use the heuristic above), equating to that of Algorithm 1. Additionally, given that the path generated by Algorithm 5 will always yield a higher score compared to that from Algorithm 1, all the theoretical results in Section 2.7 are applicable to Algorithm 5 as well.

## 3 Experimental results

We construct the genome graph with structural variations from the K-562 cancer cell line reported by Zhou et al. [2019] and the linear reference genome GRCh37, against which the SVs were called. We primarily use the VG toolkit [Garrison et al., 2018] to incorporate simple variants and further process the variant file and the resulting graph so that the final genome graph is a directed acyclic graph. Details on the construction of the genome graph can be found in Appendix H.2.

We apply our graph-based Hi-C pipeline (Section 2.1 and Appendix A) to process the raw Hi-C reads of the K-562 cancer cell line from Rao et al. [2014] (accession number: SRR1658693). Subsequently, we employed our graph-based heuristic dynamic programming algorithm to infer the sample genome. Finally, all raw Hi-C reads were remapped to the newly inferred linear genome to generate chromosome-specific contact matrices with bin size 10kb. Detailed descriptions of our algorithmic implementations, including hyper-parameter configurations, are provided in Appendix H.3.

### 3.1 Using the genome graph reference improves Hi-C read mapping

We evaluate the quality of read mapping by the number of reads from the K-562 sample SRR1658693 mapped and total number of reads that mapped perfectly without mismatches on three genome representations (Table 1). For a fair comparison between number of reads mapped to each genome, we use `vg map` [Garrison et al., 2018] for all read aligning steps, including mapping reads to GRCh37, the genome graph, and the inferred linear genome. This is because existing Hi-C read aligners such as HiC-Pro does not support sequence-to-graph alignments.

**Table 1:** Mapping statistics of Hi-C reads being mapped onto different references, computed by `vg stats -a`. Graph: reads mapped onto the genome graph; Linear reference: reads mapped onto the linear reference genome; Reconstruction: reads mapped onto the inferred linear genome. The total Hi-C reads of sample SRR1658693 is 913, 515, 598.

| SRR1658693 |  |  |  |
| --- | --- | --- | --- |
|  | Graph | Linear reference | Reconstruction |
| Mapped reads | 878,600,674 | 878,232,335 | 877,819,100 |
| Perfectly mapped reads | 301,671,424 | 277,917,664 | 289,248,295 |

As shown in Table 1, using the genome graph as a reference, both the number of total mapped reads and the number of perfectly mapped reads increase. The number of perfectly mapped reads to the inferred linear genome increases by 4.08% (more than 11 million read) compared to those perfectly mapped to the reference genome despite a slight decrease (0.05%) in total reads mapped. The reduction in total reads mapped to the inferred linear genome is partly due to the predominant presence of deletions among the large structural variations (SVs) incorporated in the genome graph (as shown in Table S1 in Appendix I). Because the inferred genome is haploid, reads that originates from the allele without the large deletions are lost. Nevertheless, the increase in perfectly mapped reads indicates that the inferred genome is more similar to the sample genome.

To examine the generality of our genome graph and inferred genome, we aligned Hi-C reads from a different K-562 sample (SRR1658694) to both the genome graph and the linear genome inferred from the sample SRR1658693. The number of perfectly mapped reads to the inferred genome increased by 4.16% (more than 24 million reads) compared to GRCh37 (Table S2 in Appendix I). The consistency between these results and those in Table 1 suggests that the inferred genome from one sample can effectively be applied to other samples within the same cell type.

The generality of genome graphs constructed on the K-562 sample can be extended to a different cell line. We align Hi-C reads from another chronic myeloid leukemia cell line, KBM-7, to the K-562 genome graph and infer the KBM-7 genome. The numbers of reads mapped to the genome graph and the inferred genome increase compared to reads mapped to the GRCh37 reference. The number of perfectly mapped reads to the inferred genome increases by 4.6% (more than 16 million reads) (Table S3 in Appendix I). This indicates potential shared structural variations between the K-562 and KBM-7 cell lines.

### 3.2 Graph Hi-C workflow improves Hi-C contact matrices

Compared to small SVs, large SVs exert a more pronounced effect on the 3D structure of chromosomes, such as Topologically Associating Domains (TADs). We validate the quality of the inferred genome by examining all four regions around SVs with lengths larger than 500 kilobases. We notice that they are all deletions. In Figure 2, Figure S5, Figure S6, and Figure S7, we show Hi-C contact matrices constructed from reads that are mapped to the inferred linear genome, the linear reference genome, and the genome graph.

**Figure 2.**
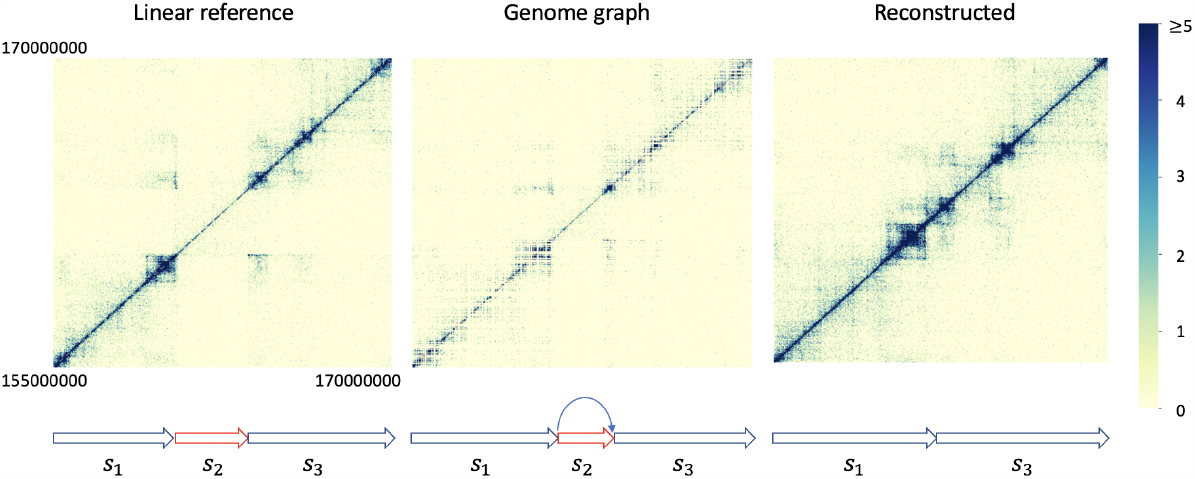
The region between 155, 000, 000bp and 170, 000, 000bp in chromosome 4. The genome graph shows a large deletion (*s*_2_) with an approximate length of 3, 000, 000bp. Note that besides this large deletion, the genome graph also contains numerous other structural variations within this region. These are not shown in the plot for the sake of clearer visualization.

The visualizations of graph-based contact matrices appear noisier compared to their counterparts due to the existence of numerous nodes that are not on the inferred path, many of which represent very short DNA sequences with few mapped reads. Consequently, these nodes generate slender stripes in contact matrices. When the reads are remapped to the inferred genome, these noisy stripes are reduced, resulting in cleaner contact matrices.

In Figure 2 and Figure S5, both the graph-based contact matrices (middle panels) and those derived using the linear reference (left panels) exhibit apparent deletion patterns, characterized by prominent stripes with very sparse interactions. These stripes also mark regions with absence of TAD structures. Our algorithm successfully identifies these deletions and produces more accurate contact matrices based on reads remapped to the inferred genome (right panels). Figure 3 presents another case where two smaller deletions of sizes 30 kb and 15 kb respectively, occur within a specific subregion of chromosome 3. Clear deletion patterns marked by stripes are evident in both the graph-based contact matrix and the matrix derived using the linear reference. Our algorithm successfully detects these two deletions, showcasing its effectiveness in identifying variations of varying sizes.

**Figure 3.**
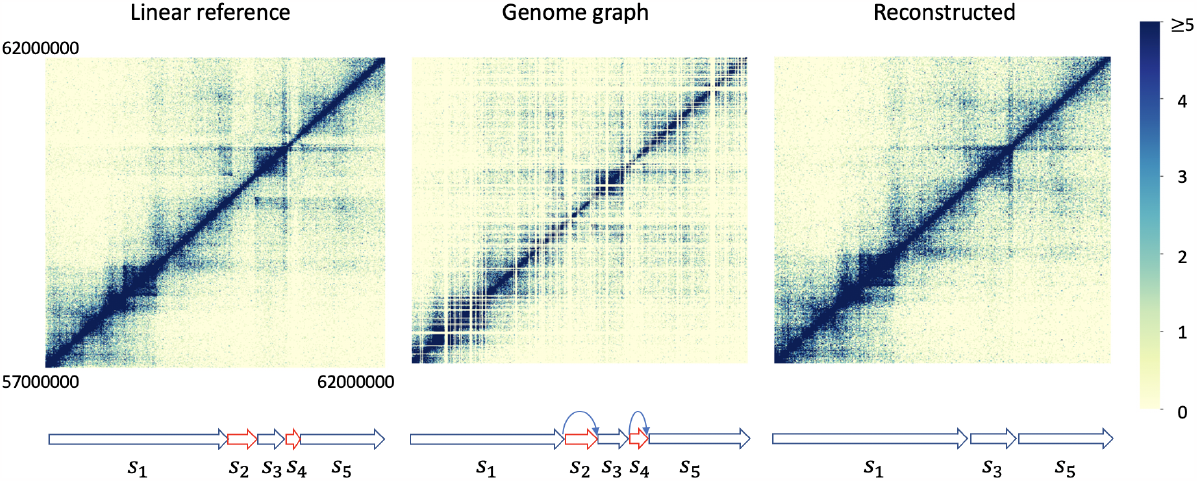
The region between 57, 000, 000bp and 62, 000, 000bp in chromosome 3. The genome graph shows two deletions (*s*_2_ and *s*_4_) with approximate lengths of 300, 000bp and 150, 000bp respectively.

In addition to incorporating true deletions in the inferred genome, we show that our graph-based Hi-C workflow is also able to discern and avoid false positive deletions in the genome graphs with few read supports (Figure S6 and Figure S7).

### 3.3 Graph Hi-C workflow improves TAD identification

We assess the quality of the new contact matrices from the inferred genome by their ability to exhibit biologically sound TAD structures. We use Armatus [Filippova et al., 2014] to identify TADs from these matrices. The detected TADs are then compared with those identified from contact matrices created from the linear reference using HiC-Pro. We evaluate the quality of TADs against the enrichment of regulatory elements CTCF and SMC in K-562 cell lines around detected TAD boundaries.

We measure the enrichment of CTCF and SMC3 around TAD boundaries with three metrics: average peak, boundary tagged ratio, and fold change (Table 2). Average peak measures the average occurrence frequency of peaks within 30 kb range centered on TAD boundaries. Boundary tagged ratio measures the frequency of TAD boundaries that are enriched for regulatory elements. Fold change measures the change of enrichment of regulatory elements between regions around and far away from TAD boundaries. Since the TAD boundaries identified using our method are situated along a path within the genome graph, we use Graph Peak Caller [Grytten et al., 2019] for calling and comparing CTCF and SMC3 peaks on the graphs. Further details on graph peak calling and these metrics can be found in Appendix H.4.

**Table 2:** The comparisons of three metrics reflecting CTCF or SMC3 enrichments near TAD boundaries across different genomes. Linear reference: linear reference genome; Reconstruction: genome inferred by our algorithm. Shortest path: genome inferred by shortest path algorithm; Longest *M* -weighted path: genome inferred by longest *M* -weighted path algorithm. TADs are called by Armatus with hyperparameter *γ* = 0.5. Hi-C sample: SRR1659693.

|  |  | Average peak | Boundary tagged ratio | Fold change |
| --- | --- | --- | --- | --- |
| CTCF | Linear reference | 0.172 | 0.346 | 0.019 |
|  | Reconstruction | **0.202** | **0.387** | **0.144** |
|  | Shortest path | 0.200 | 0.385 | 0.129 |
| | Longest $M$-weighted path | 0.199 | 0.384 | 0.139 |
| SMC3 | Linear reference | 0.091 | 0.194 | 0.044 |
|  | Reconstruction | **0.115** | **0.237** | **0.249** |
|  | Shortest path | 0.113 | 0.234 | 0.221 |
| | Longest $M$-weighted path | 0.113 | 0.233 | 0.229 |

Both CTCF and SMC3 peaks are more concentrated around TAD boundaries identified based on the inferred genomes than linear reference. Figure 4 graphically presents these peak signals around TAD boundaries, clearly showing that the signals from the inferred linear genome are more pronounced than those from the linear reference. Table 2 shows that, relative to the linear reference, there is a higher enrichment of CTCF and SMC3 signals near the TAD boundaries identified from the new contact matrices. This enhancement is robust to hyper-parameter changes during TAD calling as shown in Table S4.

**Figure 4.**
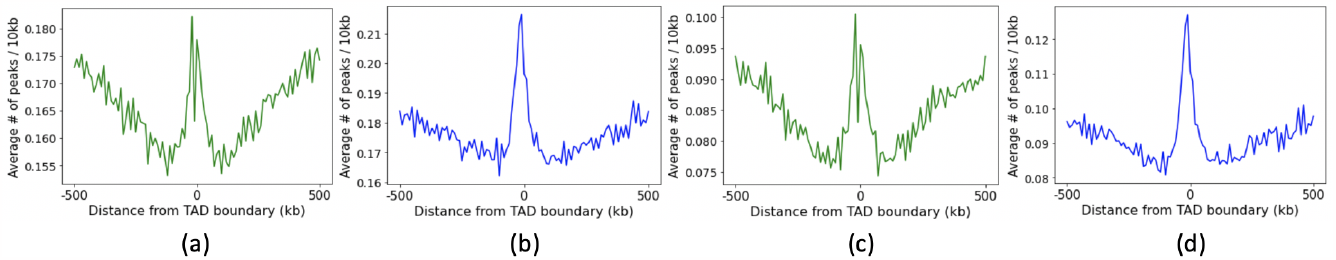
(a),(b) CTCF peak signals around TAD boundaries from the linear reference genome (a) and the inferred linear genome (b). (c),(d) SMC3 peak signals around TAD boundaries from the linear reference genome (c) and the inferred linear genome (d). TADs are called by Armatus with hyperparameter *γ* = 0.5.

To investigate the changes in TADs surrounding large structural variations, we visualize the differences between TADs identified on Hi-C matrices based on the linear reference and the inferred sample genome, shown in Figures S8, S9, and S10, which correspond to regions shown in Figures 2, S5, and 3. In the shown regions around large deletions, new TADs across the breakpoints of deletions are identified in the inferred linear genome. Additionally, very small TADs predicted within the deletion regions from the linear reference, which may not represent true TADs, are omitted in the inferred linear genome.

The improvement in TAD identification is generalizable to other K-562 samples. We evaluate TADs using new contact matrices based on Hi-C reads from sample SRR1658694 to our inferred genome from SRR1658693 (Table S5). We can draw similar conclusions that the TADs based on the inferred genome reach a higher agreement with CTCF and SCM3 elements.

We also investigate the enrichment of CTCF and SMC3 at TAD boundaries as directly inferred by our dynamic programming algorithm without remapping to the inferred genome (Table S6). The identified TADs achieve good performance in terms of average peak and boundary tagged ratio compared to TADs inferred on the linear reference genome in Table 2, which indicates that our dynamic programming algorithm can infer reasonable TAD boundaries directly from graphs.

All these findings suggest that our algorithm can successfully infer a better linear genome and generate contact matrices that more effectively capture TAD structures.

### 3.4 Dynamic programming heuristics outperforms baseline heuristics

To further assess our algorithm’s effectiveness, we compared it with two baseline approaches:

- Shortest path: This method identifies an *s*-*t* path within the genome graph, aiming for the shortest DNA sequence length, and adopts this as the inferred linear genome.
- Longest *M* -weighted path: Here, each node’s weight is determined by the total number of contacts it includes. The algorithm selects an *s*-*t* path that maximizes the sum of these weights, using the resultant path as the inferred linear genome.

The shortest path heuristic tends to be overly aggressive in handling deletions, which results in removing false positive deletions that affect a large region such as the deletion shown in Figure S7. Consequently, the resulting inferred genome is significantly shorter than the one inferred by our algorithm and therefore fewer reads are mapped (Table S7 and S8).

In contrast, the longest *M* -weighted path algorithm, despite mapping a slightly higher number of reads compared to the inferred genome by our algorithm (Table S7 and S8), lacks sensitivity to deletions and fails to identify all the major deletions as shown in Figure S11.

Furthermore, the TAD structures derived using the baseline heuristics, as reported in Tables 2 and S9, are generally less accurate compared to those generated by our dynamic programming algorithm as evaluated based on the enrichment of regulatory elements.

## 4 Discussion

In this study, we establish a novel connection between Hi-C analysis and genome graphs and explore a novel application domain in pan-genomics. We develop the first algorithm that leverages genome graphs for inferring genome sequences from Hi-C reads and the first graph-based Hi-C processing workflow which enable the efficient construction of graph-based Hi-C contact matrices. Our experimental results demonstrate that the genomes inferred via our algorithm facilitate the creation of superior Hi-C contact matrices compared to those derived using a linear reference. These promising outcomes highlight the ability of genome graphs to enhance Hi-C analysis, especially for cancer samples that contain large-scale structural variations.

There are several avenues for future research stemming from this work. First, our dynamic programming algorithm, despite its reliance on heuristics, is not exceptionally fast. For instance, processing chromosome 1 with our algorithm requires around two days, even with some parallelism techniques applied. Accelerating our algorithm could be a fruitful area of exploration. Second, currently the normalization method for Hi-C data mapped onto graphs is lacking, which is crucial for correcting inherent biases. As a result, to ensure equitable comparisons, all contact matrices presented in the experimental section of this work are unnormalized. Developing new methodologies for normalizing graph-based Hi-C data could be a vital and intriguing direction for future research. Third, our current approach, as well as that of Graph Peak Caller, is applicable only to DAGs. This limitation prevents us from testing these methods on more complex non-directed acyclic graphs, such as the human pangenome graphs created by Liao et al. [2023]. Therefore, adapting our methodology for use with general graphs represents a significant and necessary direction for future research. Additionally, given that our algorithm is applicable not only to cancer cell lines, it would be interesting to test it on more cell types, particularly normal ones, to evaluate its performance.

Finally, while there has been research like that by Wang et al. [2021] focusing on identifying structural variations from Hi-C data and rearranging contact matrices accordingly, we choose not to benchmark our method against theirs in this work. This is because our primary aim in this work is to introduce the use of genome graphs in Hi-C analysis for the first time, while the method of Wang et al., although it can create improved Hi-C contact matrices, is not able to be used on genome graphs. In the future, it would be interesting to explore the integration of these two approaches, potentially leading to even more substantial improvements in Hi-C analysis.

## 4.0.1 Acknowledgements

This work was supported in part by the US National Science Foundation [DBI-1937540, III-2232121], the US National Institutes of Health [R01HG012470] and by the generosity of Eric and Wendy Schmidt by recommendation of the Schmidt Futures program. Conflict of Interest: C.K. is a co-founder of Ocean Genomics, Inc.

## A Details of the graph-based Hi-C processing pipeline

As we describe in the main text, our Hi-C pipeline is composed of four steps. First, it constructs directed acyclic genome graphs either from various DNA sequences or from the linear reference genome coupled with Variant Call Format (`.vcf`) files. In this framework, each node of a genome graph represents a DNA sequence. Second, the pipeline performs graph-based alignment to map Hi-C reads onto the paths of the genome graphs. Following this, we prune and contract the resulting genome graph by removing paths with no reads mapped and contracting nodes into larger genomic bins to improve the computational efficiency of the following steps. Subsequently, the pipeline builds the contact matrix *M* based on the pruned graphs. Here, each dimension of the matrix represents nodes of the pruned graphs, and each matrix entry indicates the number of read pairs of which the ends are mapped to the respective nodes. There are existing approaches that can complete the first two steps. Specifically, we use the vg toolkit [Garrison et al., 2018] to construct genome graphs and facilitate the graph-based alignment. In section A.1, we introduce our approach for pruning genome graphs.

### A.1 Genome graph pruning

The genome graph constructed from vg toolkit contains unnecessary information—essentially, genetic variations that are not present in the sample from which our Hi-C data is derived. This redundant information, manifested as a subset of nodes and edges in the graph, considerably amplifies the memory usage during the execution of our Hi-C pipeline, and diminishes the computational efficiency of processes such as graph-based contact matrix construction and new genome inference. To mitigate this, we develop a novel algorithm to prune the genome graph, eliminating unnecessary nodes and edges while retaining as many useful nodes and edges as possible.

The underlying principles of our pruning algorithm are twofold: first, nodes that have no mapped Hi-C reads are more likely to be irrelevant; second, small structural variations, such as single nucleotide polymorphisms (SNPs), are unlikely to influence the 3D chromosomal structures, making their removal reasonable.

#### Algorithm 2 Genome graph pruning

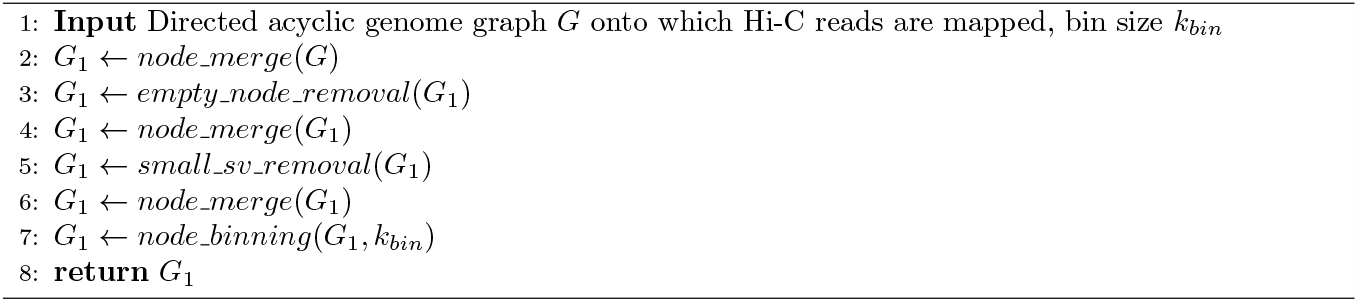

**Figure S1:**
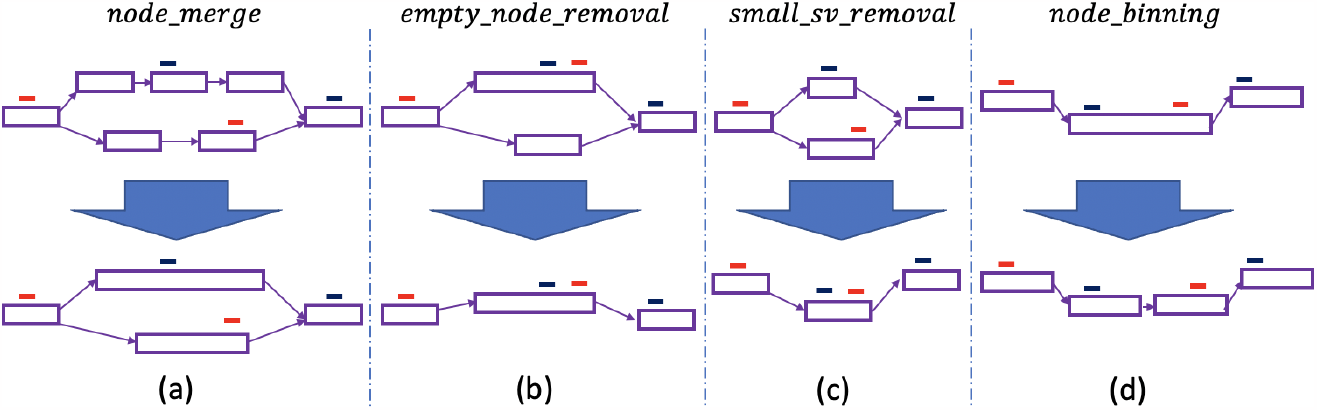
Four components of our graph pruning algorithm. Each red line represents one end of a read pair, and each blue line represents the other end.

Algorithm 2 illustrates the comprehensive framework of the graph pruning process. The algorithm takes a connected, directed acyclic genome graph *G*, onto which Hi-C reads are mapped, as its input. Leveraging the directed acyclic property of the graph will simplify the pruning procedure as well as subsequent stages in the pipeline. Section H.2 describes how we handle non-DAGs in practice. As shown in Figure S1, the pruning algorithm has four key components:

- Function *node_merge*(*G*). The indexing process of vg toolkit necessitates that each node in the genome graph represents a *k*-mer (sometimes it may represent an *s*-mer where *s < k*) [Sirén, 2017]. Typically, the value assigned to *k* is relatively small (e.g., 32), which implies that the genome graph constructed by vg toolkit has a large number of nodes, each representing a very short DNA sequence. The substantial node size will significantly impede the computational efficiency of our pipeline. To mitigate this, we borrow the idea of unitig construction, and design the function *node_merge*, which merges smaller nodes into larger ones, as illustrated in Figure S1(a). Specifically, we define two nodes, *n*_*A*_ and *n*_*B*_, as mergeable if *n*_*B*_ is the only child node of *n*_*A*_ and *n*_*A*_ is the only parent node of *n*_*B*_ (or vice versa). These two nodes can be amalgamated to create a new, larger node, representing a sequence that is the concatenation of the sequences of *n*_*A*_ and *n*_*B*_. The function *node_merge*(*G*) iteratively combines pairs of mergeable nodes until no further mergeable pairs remain. We additionally use a read projection process to project reads, which are aligned to the former nodes, onto their corresponding new nodes.
- Function *empty_node_removal*(*G*). In this function, we add a source node *s* and a sink node *t* into the DAG *G*. We also add edges from *s* to any nodes in *G* lacking a parent node, and add edges from any nodes in *G* that do not have a child node to *t*. We define a node as removable if it satisfies the following criteria: (a) it is neither the source nor the sink node, (b) it has no reads mapped onto it, and (c) each of its parent nodes has more than one child node, and each of its child nodes has more than one parent node. This third criterion ensures that the removal of this node will not significantly alter the graph’s structure. Specifically, after removing such a node, there still exists a path from *s* to every other node in *G*, and there still exists a path from any node in *G* to *t*. As shown in Figure S1(b), the function *empty_node_removal*(*G*) iteratively removes all removable nodes until no further removable nodes remain.
- Function *small_sv_removal*(*G*). We further contract our graph by removing small variants in the graph. Small variants such as SNPs and insertions and deletions ≤ 10 bases are usually represented by bubble structures [Onodera et al., 2013]. We first detect bubbles structures such that the total number of characters represented by nodes on the longest path within each bubble structure is fewer than 10 bases. Starting from the inner-most bubbles, we remove nodes that are not on the longest path within each bubble. Then, non-branching paths are merged. Reads that are mapped to the removed nodes will be mapped to the new path. The bubbles are detected using the algorithm described in Onodera et al. [2013].
- Function *node_binning*(*G, k*_*bin*_). In conventional Hi-C analysis pipelines, the aligned Hi-C reads are binned into fixed-sized genomic intervals, such as 10 kilobases, to aggregate data and smooth out noise [Lajoie et al., 2015]. Similarly, our pipeline incorporates a binning procedure applied to the graph nodes, as depicted in Figure S1(d). Specifically, when the length of the DNA sequence *s* within a node exceeds the predetermined hyperparameter *k*_*bin*_, which denotes the bin size, the *node binning* function divides the node into 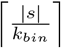 nodes. In this division, each node except for the final one represents a DNA sequence spanning a length of *k*_*bin*_ (nodes with the length of the DNA sequences smaller than *k*_*bin*_ remain unchanged). Similar to the functions *node_merge* and *small_sv_removal*, the read projection process is used here to project reads from the former nodes onto their corresponding new nodes.

Given an input DAG *G*, the graph pruning algorithm initiates by merging small nodes (line 2 of Algorithm 2). Following this, it proceeds to eliminate nodes identified as removable (line 3 of Algorithm 2). This step potentially gives rise to new mergeable nodes, prompting the algorithm to invoke the *node_merge* function once again (line 4 of Algorithm 2). Subsequently, the algorithm addresses the removal of minor structural variations (line 5 of Algorithm 2), does the *node_merge* function once again (line 6 of Algorithm 2), and culminates with the execution of node binning (line 7 of Algorithm 2).

The graph pruning algorithm yields a connected DAG that is significantly reduced in size, with Hi-C reads appropriately mapped. This pruned graph is subsequently converted into a two-dimensional contact matrix. In this matrix, each row or column corresponds to a node within the graph, and each entry represents the number of mapped read pairs between two nodes, as demonstrated in step 4 of Figure 1.

**Figure S2:**
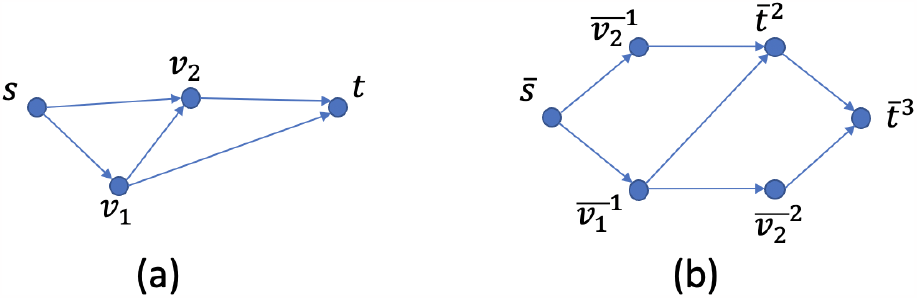
An example of converting *G* (a) to *G*^′^ (b) in the proof of Theorem 1.

## B Proof of Theorem 1

Here, we provide the proof of Theorem 1.

*Proof*. Gabow et al. [1976] proves that the path avoiding forbidden pairs problem (PAFP), introduced in Problem 2, is NP-complete in directed acyclic graphs. We now reduce PAFP on DAGs to Problem 1. Suppose we are given an instance of PAFP with a DAG *G* = (*V, E*), a source node *s*, and a sink node *t*. We define a symmetric cost function *c* on *G*, such that:

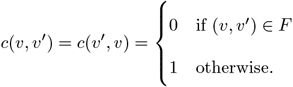

We convert *G* to a new DAG *G*^′^ = (*V* ^′^, *E* ^′^) such that every path from the source node to the sink node in the new graph has the same length (same number of nodes). We use Breadth-First-Search (BFS), starting from *s* to generate the new graph. We first create *G*^′^ that only has a source node 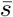, i.e. 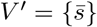and *E*^′^ = ∅. Let *V*_0_ = {*s*}. Given the node set *V*_*i*_, via BFS on *G* we create a new node set *V*_*i*+1_ which are all child nodes of the nodes in *V*_*i*_. For each node *v* ∈ *V*_*i*+1_, we add a corresponding node 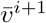 in *G* ^′^. For each node pair (*v* ^′^, *v*) such that *v*^′^ ∈ *V*_*i*_ and *v* ∈ *V*_*i*+1_, we add an edge from 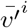 to 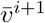 in *G*^′^ if *v*^′^ is a parent node of *v* in *G*. If 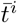 is already added in *G* ^′^, we add a node 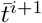 in *G*^′^ and add an edge from 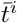 to 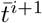. If additionally *t* is in *V*_*i*+1_, meaning that 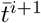 is already in *G* ^′^, we only add an edge from 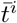 to 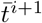 without adding the node 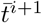 again. This procedure is iteratively conducted until *V*_*i*+1_ = {*t*} or *V*_*i*+1_ = ∅. Figure S2 shows an example of constructing *G*^′^ (Figure S2(b)) from the original graph *G* (Figure S2(a)). Since |*V*_*i*_| = *O*(|*V* |), we have that |*V* ^′^ | = *O*(|*V* |^3^). Therefore, the construction of *G*^′^ can be accomplished within polynomial time. In addition, it is easy to see that *G*^′^ is a DAG. Let 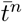 be the node in *G*^′^ that corresponds to the sink node in *G*, which was added during the final iteration of the procedure. The set of *s*-*t* paths in *G* has a one-to-one correspondence with the set of paths from 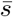 and 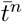 in *G* ^′^. Moreover, all the paths from 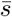 to 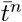 in *G*^′^ maintain equal lengths *n* + 1. We define a symmetric cost function *c*^′^ on *G* ^′^, such that:

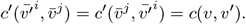

Therefore, the instance of PAFP is a yes-instance if only if there exists a 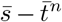 path in *G*^′^ such that the cost *c*^′^ of any node pair in this path is 1. The DAG *G*′, combined with the source node 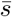, the sink node 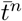, the cost function *c*′, the value *γ* = 0 and the function *μ*(*l*) ≡ 0, becomes an instance of Problem 1.

We now prove that the instance of PAFP is a yes-instance if and only if there exists a solution of Problem 1 with objective value 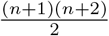, hence Problem 1 is NP-complete.

Since *γ* = 0 and *μ*(*l*) ≡ 0, the objective of Problem 1 becomes:

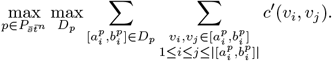

Since the cost function is non-negative, for any given 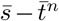 path, choosing the whole path as one domain leads to the maximal:

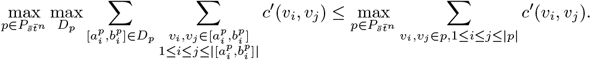

Moreover, since *c*^′^ is less than or equal to 1, and each 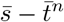 path has the same length *n* + 1, we have:

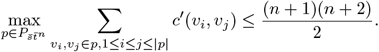

Therefore, there exists a solution of Problem 1 with objective value 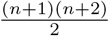 if and only if there exists a 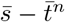 path in *G*^′^ such that the cost of any node pair in this path is 1, if and only if the instance of PAFP is a yes-instance.

## C Backtracking strategy of the dynamic programming algorithm

Here, we provide the pseudo-code of the backtracking strategy. Initiating from the sink node *t*, designated as the end node, we select the subpath where the starting node of the path is identified as the node that maximizes the expression on the right-hand side of Equation (4) (line 5 of Algorithm 3). Furthermore, this chosen subpath maximizes the function *f* across all paths extending from the start node to the end node (line 6 of Algorithm 3). The end node subsequently transitions to being a parent node of the start node that has the maximal *OPT* value (line 7 of Algorithm 3). This procedure is iteratively conducted until the start node becomes the source node *s*. The reconstructed path *p*_*opt*_ is the inferred genome we want.

### Algorithm 3 Dynamic program backtracking

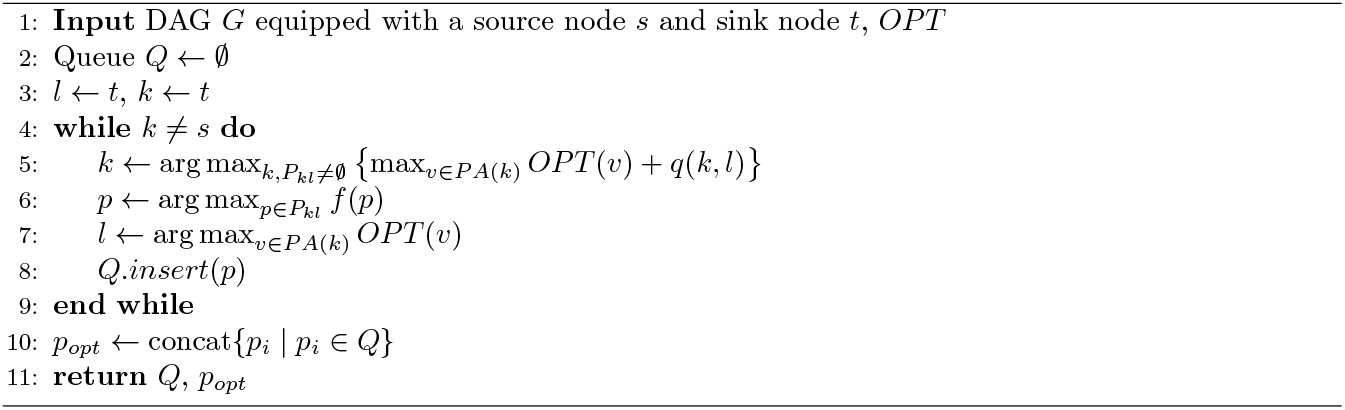

## D Proof of Proposition 1

Here, we provide the proof of Proposition 1.

*Proof*. Let *p*_*opt*_ and 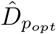be a path and its domain such that:

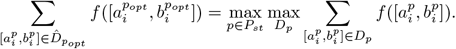

Let 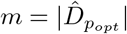, We first prove that for any *j* ≤ *m*,

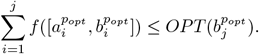

When *j* = 1, we have:

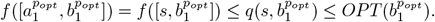

By induction,

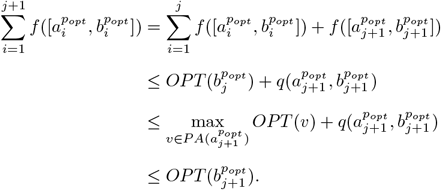

Therefore,

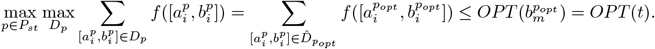

On the other hand, let us denote the output *Q* of Algorithm 3 as *Q* := {*p*_1_, *p*_2_, …, *p*_*n*_}. It is easy to see that the concatenation of all subpaths in *Q* results in forming a coherent *s*-*t* path in *G*, so we have:

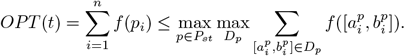

Therefore, we have 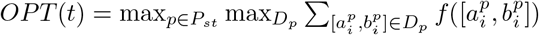. □

## E Discussion of the NP-hardness of computing the function *q*

In this section, we discuss the NP-completeness of computing function *q*. We first show that with the definition of *f* in (2), computing the function *q* in (5) is NP-complete.

### Problem 3.

Suppose we are given a directed acyclic graph *G* = (*V, E*) with a source node *s* and a sink node *t*, a pre-computed function *μ* : ℕ → ℝ_≥0_, a float value *γ* ≥ 0, and a cost function *c* : *V* × *V* → ℝ_≥0_ that maps every pair of nodes to a non-negative cost. *c* is symmetric in a sense that *c*(*v, v* ^′^) = *c*(*v* ^′^;, *v*). The problem is to find a *s*-*t* path *p* = {*v*_1_, *v*_2_, …, *v*|*p*|} over all *s*-*t* paths that can maximize the form 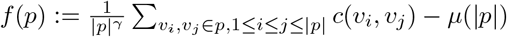, where *v*_1_ = *s* and *v*_|*p*|_ = *t* and ∀*i*, (*v*_*i*_, *v*_*i*+1_) ∈ *E*.

Since a subgraph of a directed acyclic graph is also directed acyclic, in our case of computing *q*(*k, l*), the source node is *k*, the sink node is *l*, and the cost function *c*(*v*_*i*_, *v*_*j*_) is equivalent to *M* (*v*_*i*_, *v*_*j*_).

### Theorem 3.

Problem 3 is NP-complete.

To prove Theorem 3, we first prove the NP-completeness of another problem, Problem 4, with a simpler definition of the function *f* .

### Problem 4.

We are given a directed acyclic graph *G* = (*V, E*) with a source node *s* and a sink node *t*, and a cost function *c* : *V* × *V* → ℝ_≥0_ that maps every pair of nodes to a non-negative cost. *c* is symmetric in a sense that *c*(*v, v* ^′^) = *c*(*v* ^′^, *v*). The problem is to find a *s*-*t* path *p* = {*v*_1_, *v*_2_, …, *v* _|*p*|_ } over all *s*-*t* paths that can maximize the form 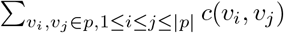, where *v*_1_ = *s* and *v*_|*p*|_ = *t* and ∀*i*, (*v*_*i*_, *v*_*i*+1_) ∈ *E*.

### Lemma 1.

Problem 4 is NP-complete.

*Proof*. The proof is highly similar to the proof of Theorem 1. We reduce PAFP on DAGs to Problem 4. Given an instance of PAFP with a DAG *G* = (*V, E*), a source node *s*, and a sink node *t*. We define a symmetric cost function *c* on *G*, such that:

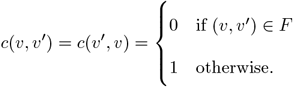

We convert *G* to a new DAG *G*^′^ = (*V* ^′^, *E* ^′^) such that every path from the source node to the sink node in the new graph has the same length (same number of nodes). We use Breadth-First-Search (BFS), starting from *s* to generate the new graph. We first create *G*^′^ that only has a source node 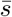, i.e. 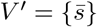 and *E*^′^ = ∅. Let *V*_0_ = {*s*}. Given the node set *V*_*i*_, via BFS on *G* we create a new node set *V*_*i*+1_ which are all child nodes of the nodes in *V*_*i*_. For each node *v* ∈ *V*_*i*+1_, we add a corresponding node 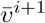 in *G* ^′^. For each node pair (*v* ^′^, *v*) such that *v*^′^ ∈ *V*_*i*_ and *v* ∈ *V*, we add an edge from 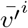 to 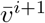in *G*^′^ if *v*^′^ is a parent node of *v* in *G*. If 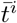 is already added in *G*′, we add a node 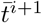 in *G*^′^ and add an edge from 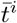to 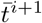. If additionally *t* is in *V*_*i*+1_, meaning that 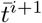 is already in *G* ^′^, we only add an edge from 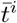 to 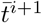 without adding the node 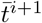 again. This procedure is iteratively conducted until *V*_*i*+1_ = {*t*} or *V*_*i*+1_ = ∅. Figure S2 shows an example of constructing *G*^′^ (Figure S2(b)) from the original graph *G* (Figure S2(a)). Since |*V*_*i*_| = *O*(|*V* |), we have that |*V* ^′^ | = *O*(|*V* |^3^). Therefore, the construction of *G*^′^ can be accomplished within polynomial time. Besides, it is easy to see that *G*^′^ is a DAG. Let 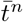 be the node in *G*^′^ that corresponds to the sink node in *G*, which was added during the final iteration of the procedure. The set of *s*-*t* paths in *G* has a one-to-one correspondence with the set of paths from 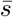 and 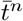 in *G* ^′^. Moreover, all the paths from 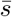 to 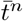 in *G*^′^ maintain equal lengths *n* + 1. We define a symmetric cost function *c*^′^ on *G* ^′^, such that:

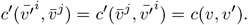

Therefore, the instance of PAFP is a yes-instance if only if there exists a 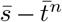 path in *G*^′^ such that the cost *c*^′^ of any node pair in this path is 1. The DAG *G* ^′^, combined with the source node 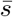, the sink node 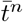, and the cost function *c* ^′^, becomes an instance of Problem 4. There exists a solution of Problem 4 with objective value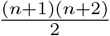 if and only if there exists a 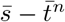 path in *G*^′^ such that the cost of any node pair in this path is 1, if and only if the instance of PAFP is a yes-instance. Therefore, Problem 4 is NP-complete. □

We now prove Theorem 3.

*Proof of Theorem 3*. Using the same strategy as the proof of Lemma 1, we convert an instance of PAFP to a new DAG *G*^′^ with the source node 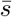, the sink node 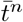, the cost function *c* ^′^, and arbitrary *γ* and *μ*, which also becomes an instance of Problem 3. Since all the paths from 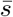 to 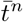 in *G*^′^ maintain equal lengths *n* + 1, if and only if the instance of PAFP is a yes-instance there exists a solution of Problem 3 with objective value 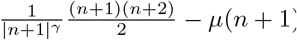. Therefore, Problem 3 is NP-complete.

The NP-completeness of *q* function computation is not attributed to the exclusive definition of function *f* as outlined in (2). We now consider some other definitions of *f* and prove that computing *q* remains NP-complete under these definitions. First, we define *f* as 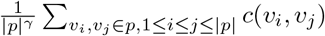 with-out *μ* function.

### Problem 5.

Given a directed acyclic graph *G* = (*V, E*) with a source node *s* and a sink node *t*, a float value *γ* ≥ 0, and a cost function *c* : *V* × *V* → ℝ_≥0_ that maps every pair of nodes to a non-negative cost. *c* is symmetric in a sense that *c*(*v, v*′) = *c*(*v*′, *v*), the problem is to find a *s*-*t* path *p* = {*v*_1_, *v*_2_, …, *v*_|*P*_ |} over all *s*-*t* paths that can maximize the form 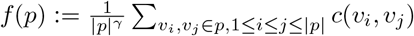, where *v*_1_ = *s* and *v*_|*p*|_ = *t* and ∀*i*, (*v*_*i*_, *v*_*i*+1_) ∈ *E*.

### Theorem 4.

Problem 5 is NP-complete.

*Proof*. Using the same strategy as the proof of Lemma 1, we convert an instance of PAFP to a new DAG *G*^′^ with the source node 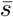, the sink node 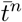, the cost function *c* ^′^, and an arbitrary *γ* ≥ 0, which also becomes an instance of Problem 5. Since all the paths from 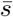 to 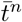 in *G*^′^ maintain equal lengths *n* + 1, if and only if the instance of PAFP is a yes-instance there exists a solution of Problem 5 with objective value 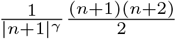 . Therefore, Problem 5 is NP-complete.

To obviate the necessity of pre-specifying the value for the hyper-parameter *γ*, we also consider the following representation of function *f*, characterized by a normalization form devoid of any pre-determined hyper-parameter:

### Problem 6.

Let us be given a directed acyclic graph *G* = (*V, E*) with a source node *s* and a sink node *t*, and a cost function *c* : *V* × *V* → ℝ_≥0_ that maps every pair of nodes to a non-negative cost. *c* is symmetric in a sense that *c*(*v, v* ^′^) = *c*(*v* ^′^, *v*). The problem is to find a *s*-*t* path *p* = {*v*_1_, *v*_2_, …, *v*_|*P*_ |} over all *s*-*t* paths that can maximize the form

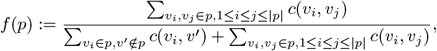

where *v*_1_ = *s* and *v*_|*p*|_ = *t* and ∀*i*, (*v*_*i*_, *v*_*i*+1_) ∈ *E*.

Here, the denominator of *f* represents the cumulative contact count between nodes within path *p* and all nodes within graph *G*, irrespective of their presence in path *p* or not. We prove in the following that, with this new definition of *f*, computing the function *q* remains NP-complete. Given the inclusion of this denominator form, the proof is significantly more intricate compared to the proofs of Theorem 3 and Theorem 4.

### Theorem 5.

Problem 6 is NP-complete.

*Proof*. A cubic graph is a graph in which all vertices have degree three. The problem of finding a maximum independent set on cubic graphs, denoted as MIS-3, is NP-complete [Choukhmane and Franco, 1986]. We prove the hardness of Problem 6 via the reduction from MIS-3.

Given an instance of a cubic graph *G*_1_ = (*V*_1_, *E*_1_), we first construct a directed acyclic graph *G*_2_ = (*V*_2_, *E*_2_). Assuming that *V*_1_ consists of *n* nodes, denoted as {*v*_1_, *v*_2_, …, *v*_*n*_} (note that the indices of these nodes can be arbitrary), we introduce two nodes in *V*_2_ for each 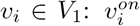 and 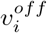 . Furthermore, we incorporate a source node *s*, a sink node *t*, and a “zombie” node *v*_0_ into *V*_2_, resulting in the set 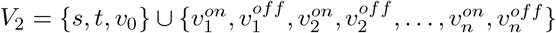.

In the new graph, we first add three directed edges from *s* to 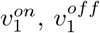, and *v*_0_. Subsequently, we add two directed edges: 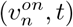 and 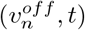. Finally, for each 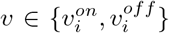, *i* ∈ {1, …, *n* − 1}, we add two directed edges 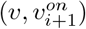 and 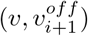. Figure S3 shows an example of constructing *G*_2_ from a cubic graph *G*_1_. It is easy to see that *G*_2_ is a DAG, and the construction can be accomplished within polynomial time.

**Figure S3:**
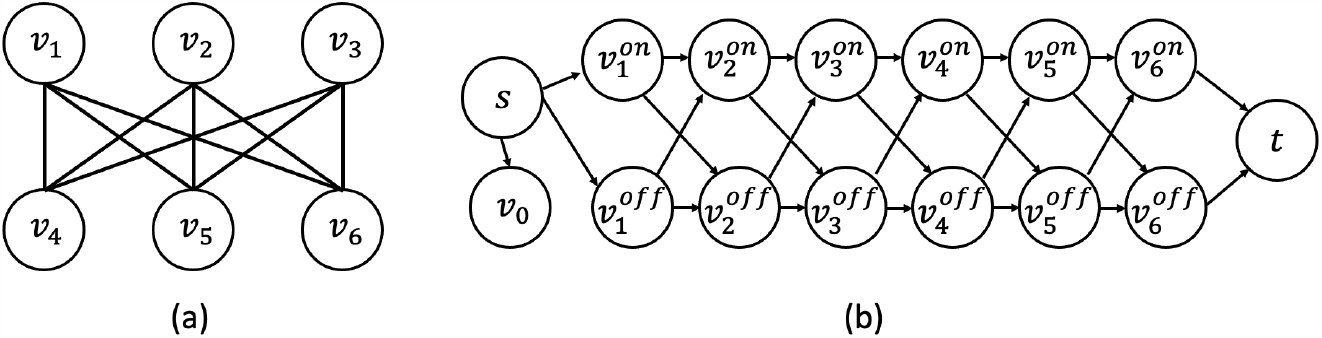
An example of generating a DAG (b) from a cubic graph instance (a) in the proof of Theorem 5.

We define the symmetric cost function *c* on *G*_2_ as the following:

- For each *v* ∈ *V*_2_, *c*(*v, v*) = 0.
- For each *v* ∈ *V*_2_, *c*(*s, v*) = *c*(*v, s*) = 0, except that 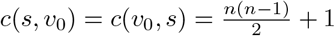.
- For each *v* ∈ *V*_2_, *c*(*t, v*) = *c*(*v, t*) = 0.
- For each *v*_*i*_, *v*_*j*_ ∈ *V*_1_, if (*v*_*i*_, *v*_*j*_) ∈ *E*_1_, then set 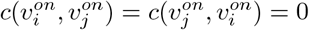
- Set 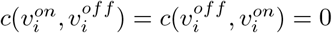 for each *i* ∈ {1, …, *n*}.
- For all the other cost values *c*(*v*_*i*_, *v*_*j*_), *v*_*i*_, *v*_*j*_ ∈ *V*_2_ that have not been defined above, set as 1.

Since

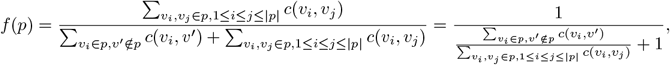

let

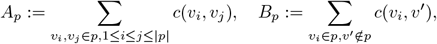

then maximizing *f* is equivalent to maximizing 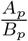.

An *s*-*t* path in *G*_2_ can be considered as a node subset selection in *G*_1_: if 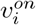 is in the path, then *v*_*i*_ ∈ *V*_1_ is selected, otherwise *v*_*i*_ is not selected. We show in the following that any path that maximizes *f* corresponds to a maximum independent set of *G*_1_.

Let *p*^∗^ be a path in *G*_2_ that maximize *f*, and define 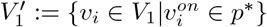. We first prove that 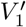 is an independent set of *G*_1_. If not, then these exists two nodes *v*_*i*_, *v*_*j*_ ∈ *V*_1_ such that (*v*_*i*_, *v*_*j*_) ∈ *E*_1_ and 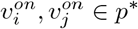. Without loss of generality, we pick *i* from {*i, j*}, and consider a new path *p* ^′^, all the nodes in *p*^′^ are the same as *p*^∗^, except that 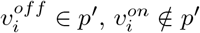. We show in the following that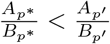:

- For the numerator *A*_*p*_, since only one node is changed 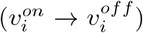, the differences between *A*_*p*_′ and *A*_*p*∗_ are all from cost values related to node *v*_*i*_. Since:
  - 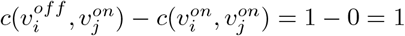
  - For any *k* ∉ {*i,j}*,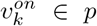, then 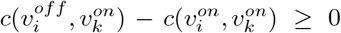; if 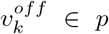 then,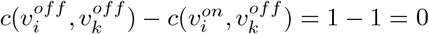;
  we have that *A*_*p*_′ ≥ *A*_*p*_∗ + 1.
- Similarly, for the denominator *B*_*p*_, the differences between *B*_*p*_′ and *B*_*p*_∗ are all from cost values related to node *v*_*i*_. Since:
  - in 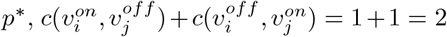 while in 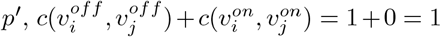, hence 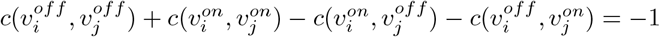,
  - For any *k* ∉ {*i,j}*,
    * if (*v*_*i*_, *v*_*k*_) ∉ *E*_1_ and 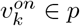,then 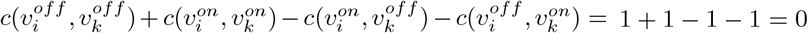,
    * if (*v*_*i*_, *v*_*k*_) ∉ *E*_1_ and 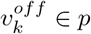,then 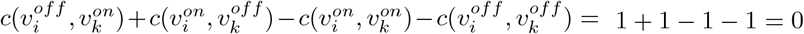,
    * if (*v*_*i*_, *v*_*k*_) ∉ *E*_1_ and 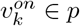,then 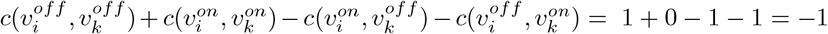,
    * if (*v*_*i*_, *v*_*k*_) ∉ *E*_1_ and 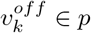,then 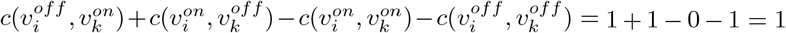 . Since *G*_1_ is a cubic graph, the number of node *v*_*k*_ that is different from *v*_*j*_ and has an edge with *v*_*i*_ is at most 2.

Therefore, *B*_*p*_′ ≤ *B*_*p*∗_ − 1 + 2 ∗ 1 = *B*_*p*∗_ + 1.

For any *s*-*t* path in *G*_2_, it is easy to see that 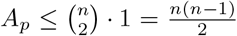, besides, 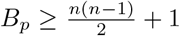 since *s* is always in the path and *v*_0_ is always not in the path, which means that *B*_*p*_ *> A*_*p*_. Therefore, we have:

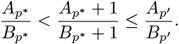

This means that *f* (*p*^′^) *> f* (*p*^∗^), leading to the contradiction. Therefore, 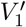 is an independent set of *G*_1_.

Next, we show that 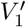 is a maximum independent set of *G*_1_. For each independent set 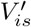 of *G*_1_, we construct a *s*-*t* path *p*_*is*_ in *G*_2_ that corresponds to it: if 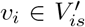, then 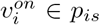, otherwise 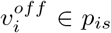.

Since 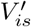 is an independent set,

- 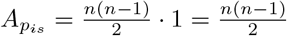
- For all those 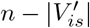 “off” nodes 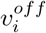 in, 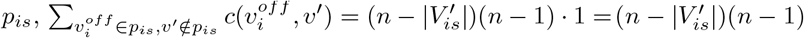, and for all those 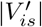 “on” nodes 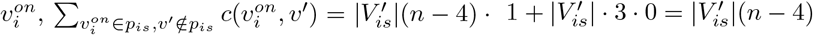. Therefore, we have:

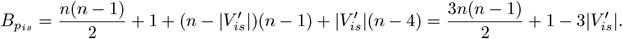

Hence:

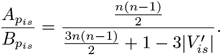

If 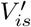 is not a maximum independent set, then by choosing a maximum independent set 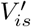 we have 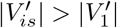. Let *p*_*is*_ be the *s*-*t* path in *G*_2_ that corresponds to 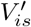, we have:

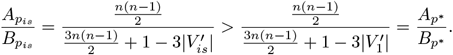

This means that *f* (*p*^′^) *> f* (*p*^∗^), leading to the contradiction. Therefore, finding an *s*-*t* path *p*^∗^ in *G*_2_ that maximizes the objective function in Problem 6 is equivalent to deciding whether there is a maximum independent set in *G*_1_ with size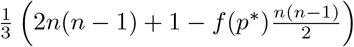. Therefore, Problem 6 is NP-complete. □

Computing the *q* function remains NP-complete if *μ* function is added into the form of *f* defined in Problem 6.

### Problem 7.

Suppose we are given a directed acyclic graph *G* = (*V, E*) with a source node *s* and a sink node *t*, a pre-computed function *μ* : ℕ → ℝ_≥0_, and a cost function *c* : *V* × *V* → ℝ_≥0_ that maps every pair of nodes to a non-negative cost. *c* is symmetric in a sense that *c*(*v, v*′) = *c*(*v*′, *v*). The problem is to find a *s*-*t* path *p* = {*v*_1_, *v*_2_, …, *v*_|*P*_ |} over all *s*-*t* paths that maximizes the form

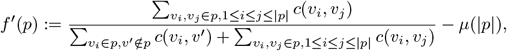

where *v*_1_ = *s* and *v*_|*p*|_ = *t* and ∀*i*, (*v*_*i*_, *v*_*i*+1_) ∈ *E*.

### Theorem 6.

Problem 7 is NP-complete.

*Proof*. Using the same strategy as the proof of Theorem 5, we convert an instance of MIS-3, *G*_1_, to a new DAG *G*_2_ with the source node *s*, sink node *t*, the cost function *c*, and an arbitrary *μ* function, which also becomes an instance of Problem 7. It is easy to see that all the paths from *s* to *t* in *G*_2_ maintain equal lengths *n* + 2, where *n* is the number of nodes in *G*_1_. Therefore, finding an *s*-*t* path *p*^∗^ in *G*_2_ that maximizes the objective function in Problem 7 is equivalent to deciding whether there is a maximum independent set in *G*_1_ with size 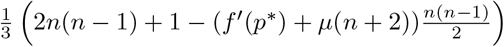 . Therefore, Problem 7 is NP-complete.

## F The NP-hardness of the function *μ* computation

According to Filippova et al. [2014], we first define the function *μ* as:

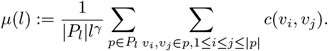

Here, *P*_*l*_ is the collection of all paths with length *l* in the directed acyclic genome graph *G* and *c*(*v*_*i*_, *v*_*j*_) is equivalent to *M* [*v*_*i*_, *v*_*j*_], the number of contacts between node *v*_*i*_ and node *v*_*j*_. With the linear reference, each node corresponds to a genomic bin. Filippova et al. [2014] demonstrated a method for efficiently pre-computing *μ* on the linear reference genome. However, while the above formulation of *μ* can be polynomially computed on a DAG, using it is not a viable option as an indicator of expected interaction frequency in our scenario. This is primarily due to the presence of unnecessary nodes within the genome graph, a phenomenon persisting even post-pruning. Consequently, we encounter redundant paths — paths not present in the sample genome from which the Hi-C data is derived. Such paths might exhibit minimal interactions, and their inclusion in the *μ* calculation would introduce significant underestimations of the expected density.

Instead, we consider another definition of *μ* on genome graphs. Let *P*_*l*_(*v*) be the collection of paths that start from node *v* and have length *l*, define *μ*(*l*) as:

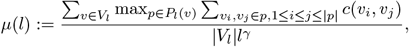

where *V*_*l*_ represents the set of nodes in the genome graph where at least one path with a length of *l* originates. For each node in the graph, we only consider the paths with maximum interactions amongst all paths originating from the same node and possessing equal length. This is because, intuitively, paths with the most interactions are more likely to represent the ground truth paths—those actually present in the sample genome. Unfortunately, we prove in the following that obtaining the value of *μ*(*l*) with the definition above for all *l* ∈ ℕ likely cannot be achieved within polynomial time.

### Theorem 7.

Let *l*_*max*_ be the length of the longest path in *G*. The problem of computing the vector 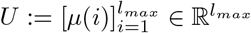 is NP-complete.

*Proof*. We prove Theorem 7 via a reduction from Problem 4. Given an instance graph *G* = (*V, E*) with source *s*, sink *t* and the cost function *c*, we assume—without loss of generality—that there exists a path from *s* to every node *v* in *G*, and a path from each node *v* to *t*. In instances where this is not the case, nodes without such paths (and their adjacent edges) can be removed within a polynomial time frame, without altering the objective value. Furthermore, we assume—without loss of generality—that *c*(*t, v*) = *c*(*v, t*) = 0 for all *v* ∈ *V* (otherwise we can add a new sink node *t*^′^ with costs *c*(*t*^′^, *v*) = *c*(*v, t*^′^) = 0 for all *v* ∈ *V* and add an edge from *t* to *t*^′^).

We use the same way to convert *G* to a new DAG *G*^′^ with source node 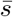 and sink node 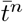. Therefore, Figure S2 is also an example of graph conversion utilized here, except that *G* in Figure S2(a) represents an instance of Problem 4 rather than a PAFP instance. We define the cost function *c*^′^ on *G*^′^ such that 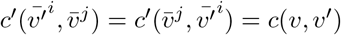.

It is easy to see that all the paths in *G*^′^ that has length *n* + 1 are from 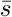 to 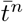, and all the paths from 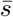 and 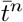 in *G*^′^ maintain equal lengths *n* + 1. Therefore, we have 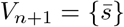 and 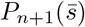 is equal to 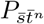, the collection of all 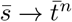 paths. Therefore, in *G*^′^,

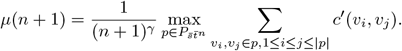

Since the set of *s*-*t* paths in *G* has a one-to-one correspondence with the set of paths from 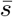 and 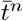 in *G*^′^, we have:

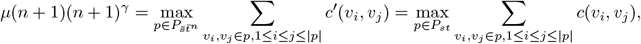

The last formula is the objective function of Problem 4. As a result, if *U* can be computed in polynomial time, then *μ*(*n* + 1) can be computed in polynomial time, which means that Problem 4 can be solved in polynomial time, leading to the contradiction. Therefore, obtaining the values of the vector 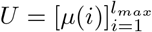 is NP-complete.

## G More discussion on the heuristic algorithm for computing *q*

### G.1 Details of the *insert* function

Here we provide the details and pseudocode of the *insert* function.

#### Algorithm 4

*insert* function

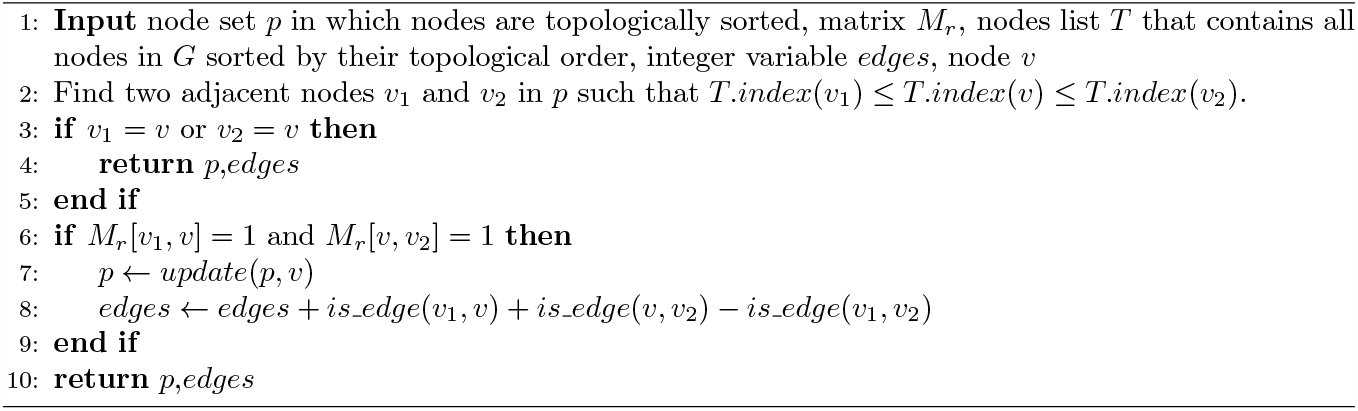

### G.2 Time complexity analysis of the heuristic algorithm

We show the proof of Theorem 2.

*Proof*. We first show that the time complexity of running Algorithm 4 once is 𝒪(log(|*V* |)) where |*V* | is the number of nodes in the graph. We use a balanced tree such as AVL tree to maintain all nodes in *p*, and use the tree to identify two adjacent nodes, *v*_1_ and *v*_2_, in *p*, where the topological ordering of *v* falls between that of *v*_1_ and *v*_2_ (line 2 of Algorithm 4). Since the existence of a path traversing all nodes within *p* has already been confirmed in earlier steps, a path that passes through all nodes in *p* ∪ *v* is existent if, and only if, there is a path from *v*_1_ to *v* as well as a path from *v* to *v*_2_ (line 6 of Algorithm 4). If the path exists, the node *v* is then inserted into the balanced tree and *p* is updated (line 7 of Algorithm 4). By using the tree structure, the time complexity of both operations, line 2 of Algorithm 4 and line 7 of Algorithm 4, is 𝒪 (log(|*p*|)). All the other operations in Algorithm 4 are 𝒪 (1). Since |*p*| ≤ |*V* |, the time complexity of Algorithm 4 is 𝒪 (log(|*V* |)).

The task of computing *f* (*p*) generally has a time complexity of 𝒪 (|*p*|^2^), as it requires summing up the interactions across all node pairs in *p*. Considering that Algorithm 1 may call the *insert* function up to |*V* | times and evaluate *f* just once, the total time complexity for Algorithm 1 is 𝒪 (|*V* | log(|*V* |) +|*V* |^2^) = 𝒪 (|*V* |^2^).

In a DAG, the reachable matrix *M*_*r*_ can be computed within 𝒪 (|*E*||*V* |) via the topological ordering, where |*E*| is the number of the edges in the graph. The dynamic program (4) only requires a single computation of *M*_*r*_ and invokes Algorithm 1 up to 𝒪 (|*V* |^2^) times. Therefore, the total time complexity of the dynamic program when utilizing the heuristic algorithm is 𝒪 (|*E*||*V* | + |*V* |^4^) = 𝒪 (|*V* |^4^).

### G.3 Worst case example

**Figure S4:**
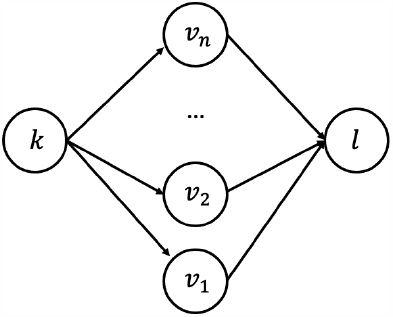
A worst-case example of the heuristic algorithm

We construct a worst-case family of instances for our heuristic algorithm. Let 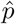 represent the path from node *k* to node *l* as predicted by Algorithm 1, and let *p*_*gt*_ denote the “ground truth” path, defined as 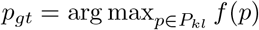. Consider the graph shown in Figure S4: there are *n* + 2 nodes in this DAG, *V* = {*k, l, v*_1_, *v*_2_, …, *v*_*n*_}. For each *i* ∈ {1, …, *n*}, there is an edge from *k* to *v*_*i*_, (*k, v*_*i*_), and an edge from *v*_*i*_ to *l*, (*v*_*i*_, *l*). We create a contact matrix *M* on this graph, defined as:

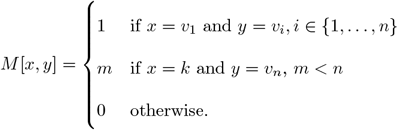

In this example, since:

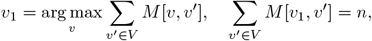

our heuristic algorithm will pick the node *v*_1_ and output the path 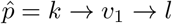. The total interactions within 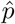 is equal to 0,

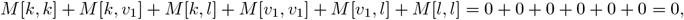

while *p*_*gt*_ is the path *k* → *v*_*n*_ → *l*, with the interactions:

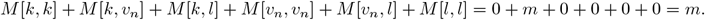

Therefore, we have:

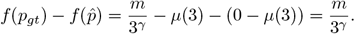

Since the values of *m* and *n* can be chosen to be arbitrarily large (as long as *m < n*), the discrepancy between 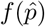 and *f* (*p*_*gt*_) can be infinitely large.

### G.4 Theoretical framework justifying the accuracy of the heuristic algorithm

As we describe in the main text, within the scope of Hi-C analysis, the distribution of interactions on a genome graph is not arbitrary. We construct a theoretical framework that more accurately reflects the real-world Hi-C situation, and demonstrate that with high probability our heuristic algorithm can output the path 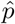 that is equivalent to *p*_*gt*_ as long as the number of mapped read pairs (interactions) is not too small.

Suppose we have a directed acyclic genome graph *G* = (*V, E*) with source node *s* and sink node *t*, and we specify a ground-truth path *p*_*gt*_ connecting *s* and *t*. Let 𝒳 denote all the ordered node pairs of on *G*: 𝒳 = {(*v*_1_, *v*_2_) | *v*_1_, *v*_2_ ∈ *V, v*_1_ ≤_*top*_ *v*_2_}, where ≤_*top*_ denotes the inequality under the topological ordering. We will define a probability distribution *P* (*X*) on *G* and assume the interaction pairs are independent and identically distributed (iid) samples from it. Specifically, we assume that the probability mass function *p*(*x*) = ℙ(*X* = *x*) of the distribution takes the following mixture form:

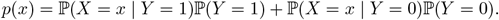

Here, *Y* is a random variable indicating from which latent state the sample is drawn: *Y* = 1 represents the “ground truth” state, while *Y* = 0 represents the “noise” state. The samples from the noise state are less informative—We assume that the interaction is sampled from a uniform distribution over 𝒳 :

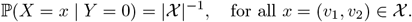

In the “ground truth” state, we assume that both of the nodes of the interaction lie on the path *p*_*gt*_. Therefore, the support of *X* conditioned on *Y* = 1 is a subset 𝒳 _*sub*_ of 𝒳, which is defined as 𝒳 _*sub*_ = {(*v*_1_, *v*_2_) | *v*_1_, *v*_2_ ∈ *p*_*gt*_, *v*_1_ ≤_*top*_ *v*_2_}. Moreover, as shown in previous work such as Ay et al. [2014], in Hi-C experiments the probability of observing an intra-chromosomal interaction between two chromosomal locations is inversely related to their genomic distance: the smaller the distance, the higher the likelihood. Formally,

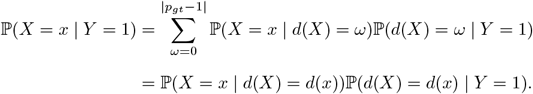

Here, *d*(*x*) = *d*((*v*_1_, *v*_2_)) is the number of edges between *v*_1_ and *v*_2_ in *p*_*gt*_, which indicates the distance between *v*_1_ and *v*_2_. Similar to Ay et al. [2014], we further assume that the probabilities of observing interactions at the same distance are equivalent. This implies that ℙ(*x*|*d*(*x*)) = {|*p*_*gt*_| − *d*(*x*)}^−1^ because the number of unique interactions with a distance value *d*_0_ in the path *p*_*gt*_ equals |*p*_*gt*_| − *d*_0_, where |*p*_*gt*_| denotes the number of nodes in *p*_*gt*_. In summary, we have the following assumption about *p*(*x*):

#### Assumption 1.

Given a DAG *G* = (*V, E*) with source node *s* and sink node *t* and the ground truth path *p*_*gt*_ connecting them, the probability mass function *p*(*x*) can be factorized as follows:

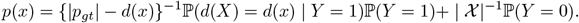

We further assume the probability of sampling from the ground truth state is strictly greater than zero: ℙ(*Y* = 1) *>* 0.

Furthermore, we have an assumption on ℙ(*d*(*X*) = *d*(*x*) | *Y* = 1):

#### Assumption 2.

From any *D > d* ≥ 0, we assume

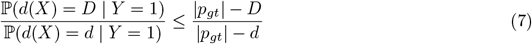

#### Assumption 2

states the probability of sampling an interaction with a larger distance is smaller than that of one with a smaller distance. Moreover, such a decay rate, as with respect to the distance value, is faster than linear. This is a mild assumption. In fact, as illustrated in Figure 1 of Ay et al. [2014], the count of intra-chromosomal interactions typically decays exponentially with an increasing genomic distance, which is much faster than our assumed linear rate. When the equality holds in (7), combined with the factorization Assumption 1, the sample probability ℙ(*X* = *x* | *Y* = 1) is also a constant: In this case, the samples are generated from uniform distributions under both the ground truth and the noise states.

In practical applications, we generally find that these two assumptions hold true while calculating *q*(*k, l*) for the majority of node pairs (*k, l*). Nonetheless, these assumptions may not apply in cases involving large deletions. As described in Section 2.8, we have implemented additional modifications to our algorithm to effectively address such scenarios.

Given a set of iid interaction pairs ℐ = {*x*_*i*_ ∈ 𝒳, *i* = 1, …, | ℐ |}, our main result on this theoretical model is as the following:

#### Theorem 8.

Under Assumptions 1 and 2, for any *k >* 0, the predicted path, 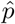, from the heuristic algorithm is identical to the ground truth path *p*_*gt*_ with a probability greater than 1 − exp(−*k*) so long as the number of interactions | ℐ | is greater than *C*|*p*_*gt*_|(log(|*V* |) + *k*). Here *C* is a constant whose value only depends on ℙ(*Y* = 1).

In Hi-C datasets, | ℐ | denotes the total number of read pairs. To prove Theorem 8, we need the following theorem, which states that the probability of 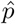deviating from *p*_*gt*_ decreases exponentially with respect to | ℐ |.

#### Theorem 9.

Given the set of interactions ℐ = {*x*_*i*_ ∈ 𝒳 } where 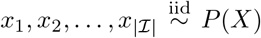. Under the Assumptions 1 and 2, we have:

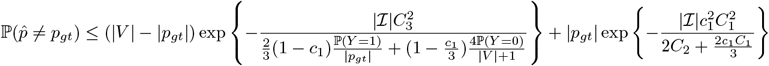

for any 0 *< c*_1_ *<* (*C*_1_|*p*_*gt*_|)^−1^ℙ(*Y* = 1). Here, *C*_1_, *C*_2_ and *C*_3_ are constants whose value does not depend on | ℐ |:

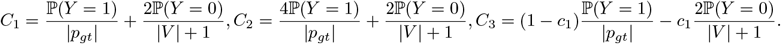

We first sketch the overarching concept behind the proof. If the output 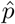 from our algorithm is different from *p*_*gt*_, then there exists a node outside of *p*_*gt*_ that is selected by the algorithm before all the nodes from *p*_*gt*_ are chosen. Let *C*(*v*) be the number of the interactions in ℐ that include node *v*, i.e., 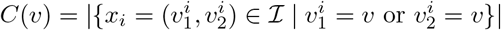. Then we have:

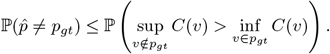

Therefore, 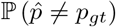 can be upper bounded if 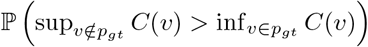 is bounded. The ideas to bound 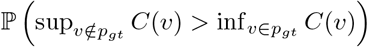 are two fold:

- By using the concentration inequalities, we show that as the value of | ℐ | increases, 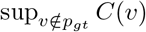 approaches a constant value *A*, while 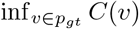 approaches another constant value *B*. These constants are correlated with the expected values respective to each.
- We show that, since *A < B*, the likelihood that 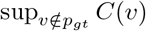 exceeds 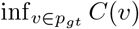diminishes as |ℐ | increases.

To prove Theorem 9, we need the following several lemmas. Recall that *C*(*v*) be the number of the interactions in ℐ that include node *v*, i.e. 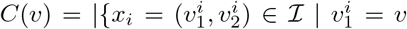or 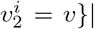. The first lemma quantifies the average number of interactions containing any fixed node on the ground truth pathway.

#### Lemma 2.

For any *v* ∈ *p*_*gt*_,

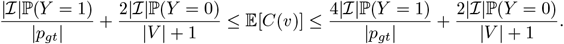

*Proof*. Let *v*_0_ be a fixed node on the ground truth path *p*_*gt*_. We define *d*_0_ := min(*d*(*s, v*_0_), *d*(*v*_0_, *t*)) ≥ 0 and *d*_1_ := max(*d*(*s, v*_0_), *d*(*v*_0_, *t*)) = |*p*_*gt*_| − 1 − *d*_0_ ≥ 0.

Consider a binary random variable *Z*_*i*_, such that:

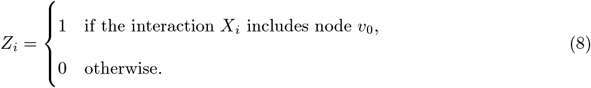

We can directly verify that 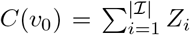. To study *E*[*C*(*v*_0_)] we just need to study *E*[*Z*_*i*_]. Since *X*_1_, *X*_2_, …, *X*_| ℐ|_ are independent random variables, we have that *Z*_1_, *Z*_2_, …, *Z*_| ℐ|_ are independent random variables as well.

In the “ground truth” state, the probability of *v*_0_ being in one sampled interaction is (law of total probability):

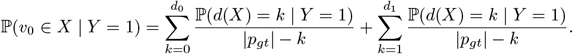

Applying Assumption 2, we have:

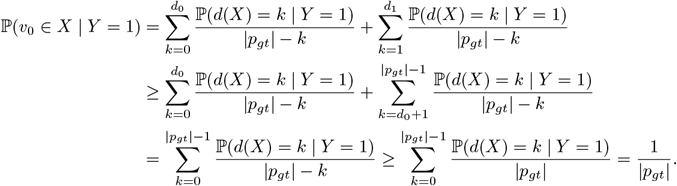

The last inequality holds because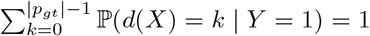. Furthermore, using Assumption 2 again, we have:

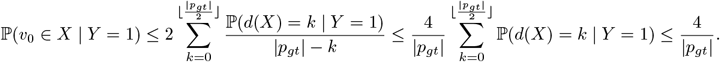

In the “noise” state, since every interaction in 𝒳 is equivalent, 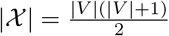, and since the number of interactions that contain *v*_0_ is |*V* |, we know that the probability of *v*_0_ being in one sampled interaction is 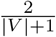

Therefore, for each *i* ∈ {1, …, |ℐ |},

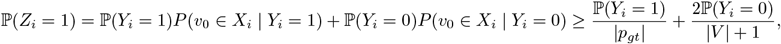

Similarly,

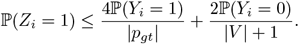

Therefore,

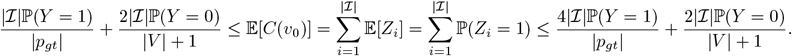

Since *v*_0_ is arbitrarily chosen, the inequalities above hold for all *v* ∈ *p*_*gt*_.

The next lemma bounds the variance of number of interactions containing a node on *p*_*gt*_.

#### Lemma 3.

For each *v* ∈ *p*_*gt*_,

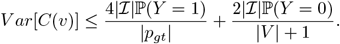

*Proof*. Let *Z*_1_, *Z*_2_, …, *Z*_|ℐ |_ denote the indicator variables defined in (8). Since they are independent, we have:

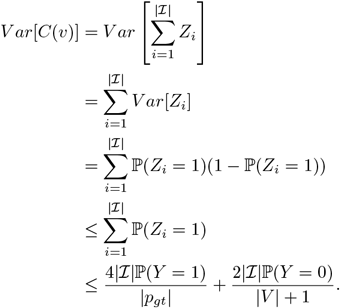

Based on Lemma 2 and Lemma 3, we can prove the following lemma, which gives a lower bound of the constant value *B*.

#### Lemma 4.

Let 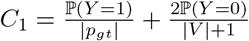 and 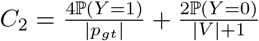, then for any 0 *< c*_1_ ≤ 1, we have

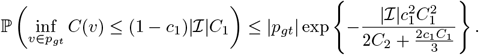

*Proof*.

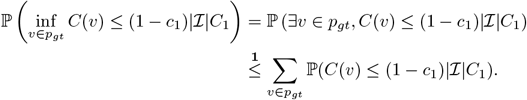

Inequality 1 holds because of the union bound. According to Bernstein inequality and the fact that for all *i*, |*Z*_*i*_| ≤ 1. We have that for all *t >* 0,

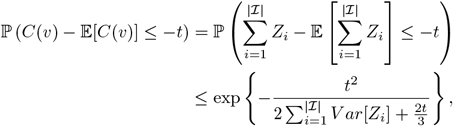

and

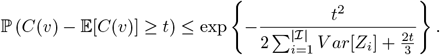

Since |ℐ |*C*_1_ ≤ 𝔼[*C*(*v*)] (Lemma 2) and 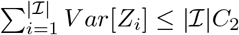 (Lemma 3), we have:

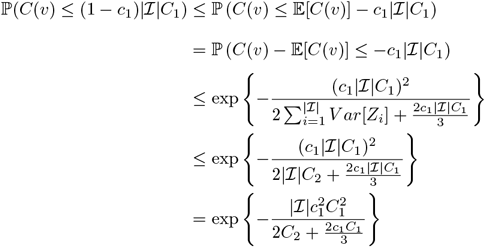

Therefore,

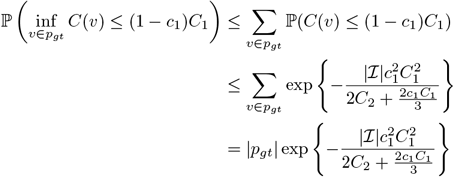

The next lemma quantifies the average number of interactions containing any fixed node not on the ground truth pathway.

#### Lemma 5.

For each *v* ∈*/ p*_*gt*_,

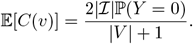

*Proof*. We use the same strategy as Lemma 2. Since nodes that are outside of *p*_*gt*_ can only be observed in interactions that are from the “noise” state, the probability of a node outside of *p*_*gt*_ being in one sampled interaction is

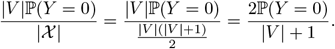

Therefore, 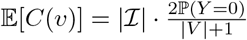 for each *v* ∈*/ p*_*gt*_.

The next lemma bounds the variance of number of interactions containing a node not on *p*_*gt*_.

#### Lemma 6.

For each *v* ∈*/ p*_*gt*_,

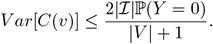

*Proof*. We use the same strategy as Lemma 3. Given an arbitrary *v* ∈*/ p*_*gt*_, consider a binary random variable *W*_*i*_ such that:

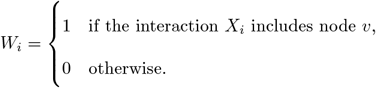

Then 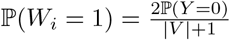 . Therefore,

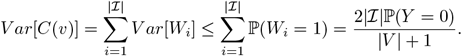

Lemma 5 and Lemma 6 will help us derive an upper bound of the constant value *A*. Now we are ready to prove Theorem 9.

*Proof of Theorem 9*. Let 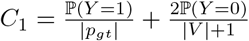, for any c_1_ such that

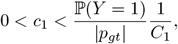

(for example, *c*_1_ can be ℙ(*Y* = 1)*/*3) we have:

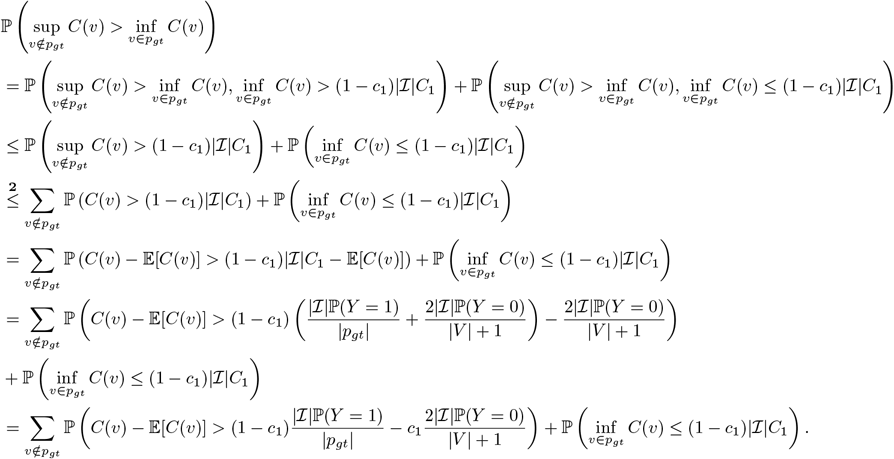

We used the union bound to get inequality 2. Let:

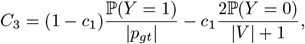

by using Bernstein inequality again, We have that for *v* ∈*/ p*_*gt*_:

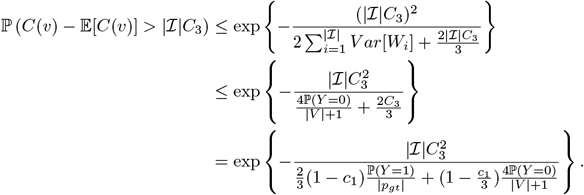

Let 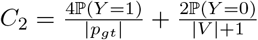, we have:

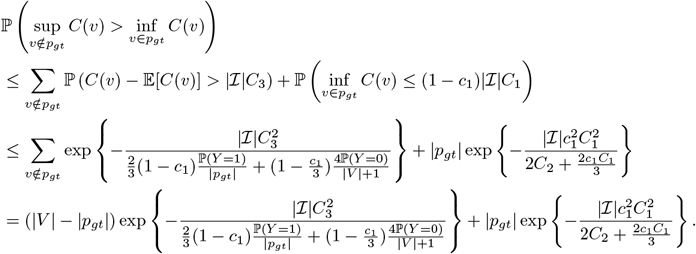

Therefore, we have:

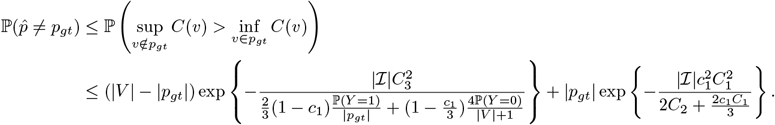

Based on Theorem 9, we can now prove Theorem 8:

*Proof of Theorem 8*. Since |*V* | ≥ |*p*_*gt*_|, we have that:

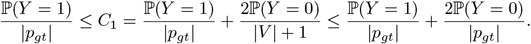

Therefore, the inequality in Theorem 9 holds for any 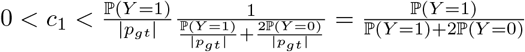,and in this region,

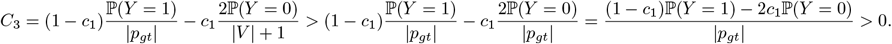

We also have:

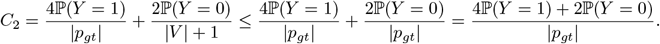

Therefore,

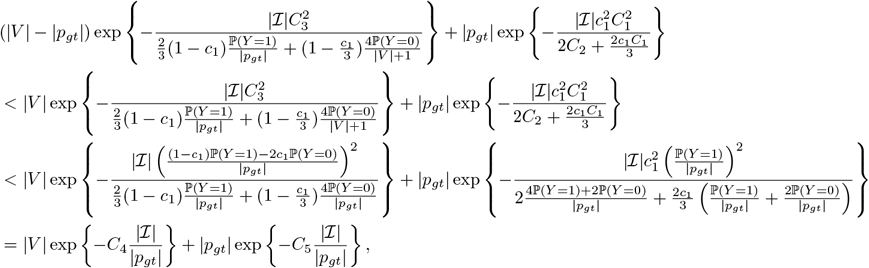

Where

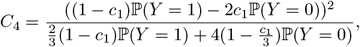

and

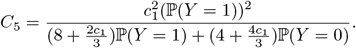

Therefore, given a small positive value *ϵ >* 0, when

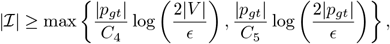

we have:

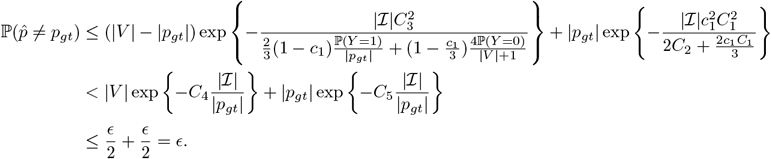

It is easy to see that, if we choose *c*_1_ to be ℙ(*Y* = 1)*/*3, and since ℙ(*Y* = 1) + ℙ(*Y* = 0) = 1, both *C*_4_ and *C*_5_ are constants whose values only depend on ℙ(*Y* = 1).

Our dynamic program computes the function *q* up to 𝒪 (|*V* |^2^) times. Therefore, to ensure a high likelihood that each predicted path matches the ground truth path, we need to establish a bound of the following form:

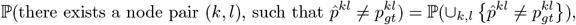

where 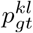 is the predicted path from node *k* to node *l* and 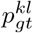 is the ground truth path from *k* to *l*. By using the union bound, we have:

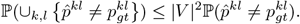

Let 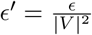, when

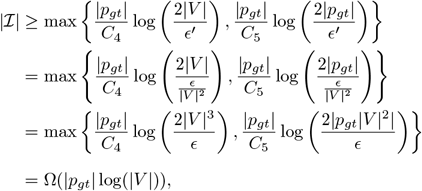

we have that

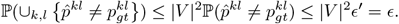

Therefore, we can see that ensuring a high likelihood of each predicted path consistent with the ground truth path doesn’t affect the sample complexity of our heuristic algorithm; it only influences the constant factor.

### G.5 Adjustment of Algorithm 1

Here, we provide the pseudo-code of Algorithm 5.

## H Implementation details of experiments

### H.1 Data availability

Raw Hi-C reads of GM12878 cell line were downloaded from Sequence Read Archive (SRA) under the accession numbers SRR1658570, SRR1658571, SRR1658572, SRR1658573, SRR1658574, SRR1658575, SRR1658576, SRR1658577, SRR1658578, SRR1658579, SRR1658580, SRR1658581, SRR1658582, SRR1658583, SRR1658584, SRR1658585, SRR1658586, SRR1658587, SRR1658588, SRR1658589, SRR1658590, SRR1658591,

#### Algorithm 5

*q*(*k, l*) computation v.2

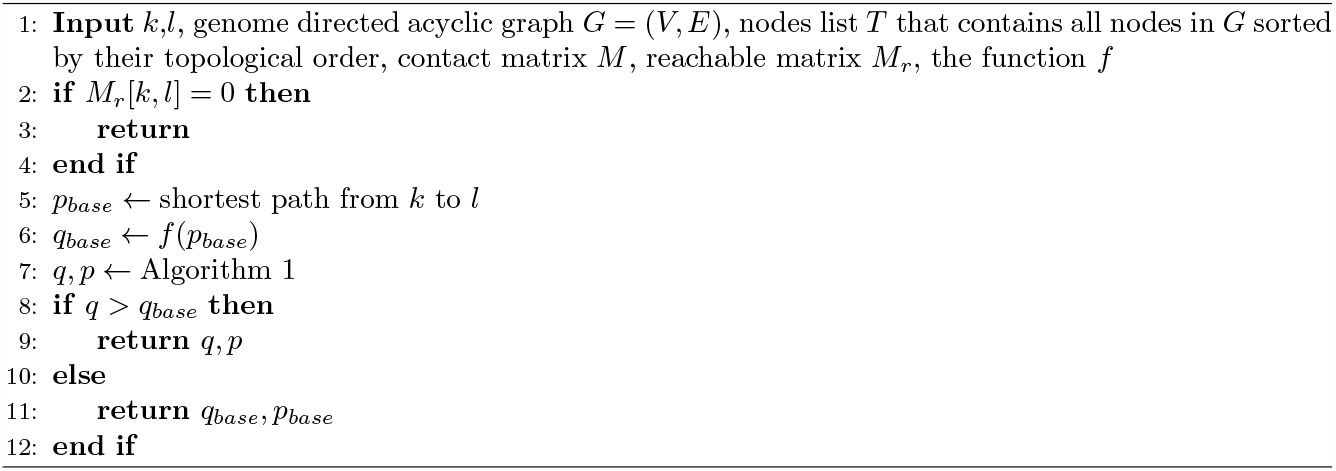

SRR1658592, SRR1658593, SRR1658594, SRR1658595, SRR1658596, SRR1658597, SRR1658598, SRR1658599, SRR1658600, SRR1658601, SRR1658602, and SRR1658603. Raw Hi-C reads of K-562 cancer cell line were downloaded from SRA under the accession numbers SRR1658693 and SRR1658694. Raw Hi-C reads of KBM-7 cancer cell line were also downloaded from SRA under the accession number SRR1658708. The `.vcf` files containing structural variations of the K-562 cell line were downloaded from ENCODE Portal (https://www.encodeproject.org) under accession numbers ENCFF356GYS, ENCFF538YDL, ENCFF574MDJ, ENCFF752OAX, ENCFF785JVR, ENCFF863MPP and ENCFF960SSF. Raw CTCF ChIP-seq reads of K-562 cell line were downloaded from ENCODE under accession numbers ENCFF001HTO and ENCFF001HTP. Raw SMC3 ChIP-seq reads of K-562 cell line were downloaded from ENCODE under accession numbers ENCFF000YZX and ENCFF000YZY.

### H.2 Genome graph construction

We use linear reference genome GRCh37 and structural variations of K-562 cancer cell line reported by Zhou et al. [2019] to construct of our genome graph via the following steps. First, seven `.vcf` files were downloaded from ENCODE Portal. These files contain variants of the K-562 cell line, categorized into three distinct types:

- Small variants. These variants includes small insertion, deletions and single-nucleotide variants that are represented by precise DNA sequence changes in the vcf, for example: REF: TCG, ALT: T.
- Large structural variants. These SVs are represented by abbreviations such as INV (inversion), DEL (deletion), INS (insertion) and DUP (duplication).
- Complex rearrangements. These SVs are represented by a set of novel adjacencies such as [chr1:6777707[G. The meanings of these symbols are described in https://samtools.github.io/hts-specs/VCFv4.2.pdf.

We then use `vg construct` command to construct genome graphs using the linear reference and `.vcf` files. Currently, `vg construct` only supports all forms of small SVs and three types of intrachromosomal large SVs: INV (inversion), DEL (deletion), and INS (insertion). We observed that certain complex rearrangements could be reformatted into recognizable abbreviations, and therefore we manually reformat them to enable their integration using `vg construct`. Furthermore, since our algorithm necessitates a directed acyclic graph (DAG) as input, we transformed non-DAG segments, specifically those resulting from inversions (INVs), into DAG-compatible structures. This transformation was done by substituting inversions with new nodes that contain the reverse complement of the DNA sequences. Although the `vg mod --unfold` function may be an alternative approach to dagify the graph, the VG team has indicated that this function is somewhat outdated and less maintained, as discussed in the GitHub issues https://github.com/vgteam/vg/issues/4103 and https://github.com/human-pangenomics/hpp\pangenome\resources/issues/22.

Table S1 presents statistics for the structural variations (SVs) ultimately integrated into our genome graph. It shows that small SVs constitute the bulk of these variations. Among the large SVs, deletions are predominant, with no instances of insertions or duplications noted.

**Table S1:** Statistics of SVs incorporated into our genome graph.

|  | small SVs | DEL | INV | other SVs |
| --- | --- | --- | --- | --- |
| Number | 3822549 | 6069 | 53 | 0 |
| Percentage | 99.84% | 0.159% | 0.001% | 0% |
| Max length (bp) | 572 | 19018917 | 365625 |  |
| Mean length (bp) | 2 | 9748 | 22127 |  |

### H.3 Implementation details of graph-based dynamic programming algorithm

We use HiC-Pro [Servant et al., 2015] to process the raw Hi-C reads of GM12878 cell line from Rao et al. [2014], and use Armatus [Filippova et al., 2014] on the contact matrices of GM12878 with bin size 10kb to compute *μ*_0_ used in Eq. (3).

In practice, as outlined in Section A, while the majority of nodes in our genome graph have sequences that are precisely 10kb in length, there exists a subset of nodes with sequences shorter than 10kb. In addition, Armatus only computes values of *μ*_0_(*l*) for every integer value of *l*. To refine the estimation of *μ*, we initially employ a spline model [Späth, 1995] for interpolation. This approach enables us to derive *μ*_0_(*l*) for every float value of *l*. Subsequently, for a given path *p*, we compute its respective *μ* value using the formula *μ*(*L*(*p*)*/*10000), where *L*(*p*) denotes the aggregate length of the DNA sequence along path *p*.

Throughout our experiments, we maintain *γ* in Eq. (2) as 0.1 and set the bin size *k*_*bin*_ to 10kb.

### H.3 ChIP-seq peak calling and analysis on graphs

We download raw CTCF and SMC3 ChIP-seq reads of the K-562 cell line from ENCODE. Following the steps described in Liao et al. [2023] (https://doi.org/10.5281/zenodo.6564396), we align these reads to our genome graph using `vg map`, and call peaks using Graph Peak Caller (v1.2.3) [Grytten et al., 2019]. To compare peaks called on the graph with TAD boundaries identified on the linear reference, we use the command `graph_peak_caller peaks_to_linear` to project all peaks to the path of the graph corresponding linear reference.

To evaluate TAD boundaries called from contact matrices derived using different genomes, we use three metrics: the average peak around TAD boundaries, the boundary tagged ratio, and the fold change. They are introduced in Zufferey et al. [2018], Liu et al. [2022] and Sefer [2022]. Since the bin size in our experiments is 10kb, we adjust the definitions of these three metrics as the following:

- Average peak around TAD boundaries, a metric that describes the density of the occurrence frequency of regulatory elements such as CTCF and SMC3 around the TAD boundaries. It is defined as:

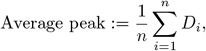

where *n* is the number of unique TAD boundaries and *D*_*i*_ is the average frequency of occurrence of regulatory elements per 10kb within a 30kb range centered on the *i*-th TAD boundary (the boundary and its two adjacent bins).
- Boundary tagged ratio, a metric that describes how frequent TAD boundaries enriched for regulatory elements are. It is defined as:

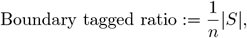

where *n* is the number of unique TAD boundaries and *S* is the set of TAD boundaries on which a regulatory element occurs within a centered 30kb range.
- Fold change, a metric that describes how much the occurrence density of regulatory elements changes between the regions near the TAD boundaries and the regions far away from the TAD boundaries. It is defined as:

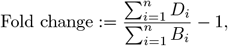

where *n* is the number of unique TAD boundaries, *D*_*i*_ is the average frequency of occurrence of regulatory elements per 10kb within a 30kb range centered on the *i*-th TAD boundary, and *B*_*i*_ is the average frequency of occurrence of regulatory elements per 10kb in bilateral regions on both sides 200kb to 500kb from the *i*-th TAD boundary.

## I Additional experimental results

We present additional experimental results.

**Table S2:** Mapping statistics of Hi-C reads being mapped onto different references, computed by `vg stats -a`. Graph: reads mapped onto the genome graph; Linear reference: reads mapped onto the linear reference genome; Reconstruction: reads mapped onto the inferred linear genome. The total Hi-C reads of sample SRR1658694 is 1, 183, 709, 106.

| SRR1658694 |  |  |  |
| --- | --- | --- | --- |
|  | Graph | Linear reference | Reconstruction |
| Mapped reads | 1,167,029,589 | 1,166,821,533 | 1,166,203,299 |
| Perfectly mapped reads | 631,181,248 | 580,395,980 | 604,569,568 |

**Table S3:**
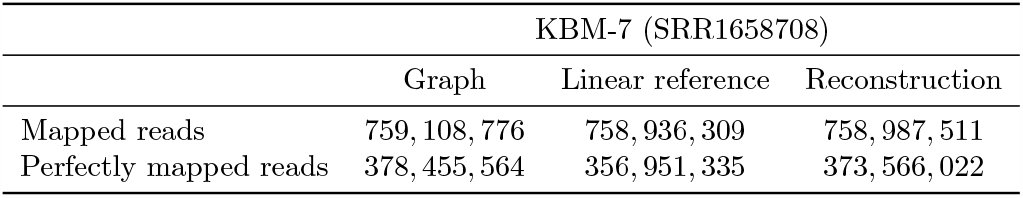
Mapping statistics of Hi-C reads being mapped onto different references, computed by `vg stats -a`. Graph: reads mapped onto the genome graph; Linear reference: reads mapped onto the linear reference genome; Reconstruction: reads mapped onto the inferred linear genome. The total Hi-C reads of sample SRR1658708 is 776, 936, 020.

**Figure S5:**
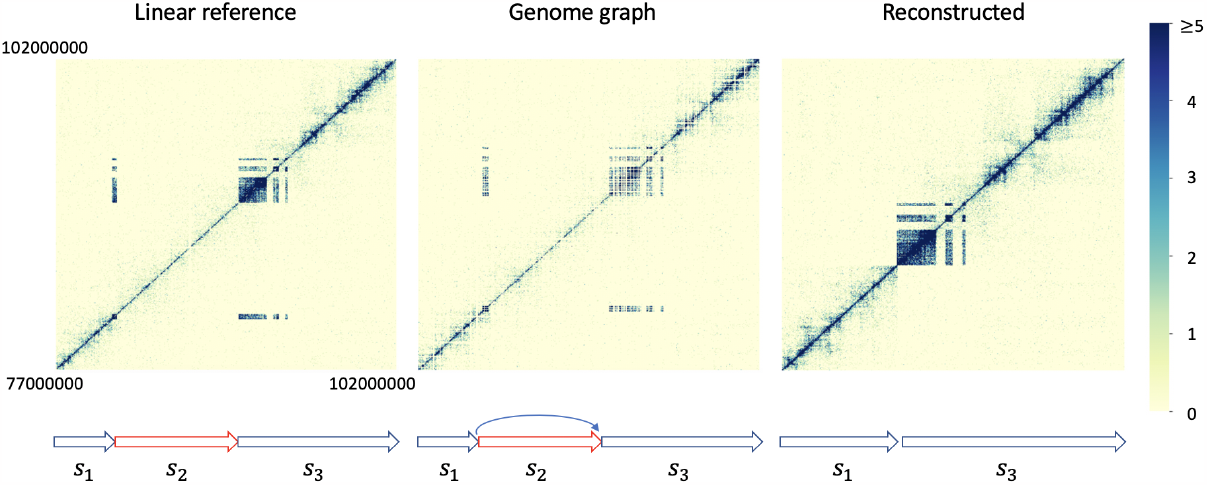
The region between 77, 000, 000bp and 102, 000, 000bp in chromosome 13. The genome graph shows a large deletion (*s*_2_) with an approximate length of 9, 000, 000bp.

**Figure S6:**
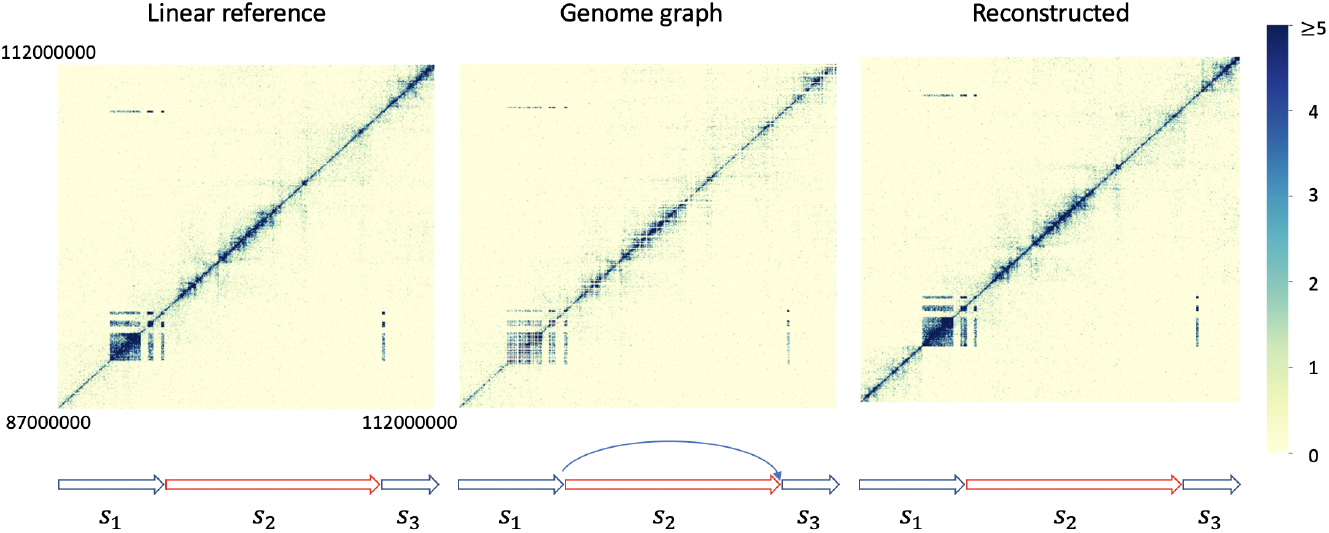
The region between 87, 000, 000bp and 112, 000, 000bp in chromosome 13. The genome graph shows a large deletion (*s*_2_) with an approximate length of 15, 000, 000bp.

**Figure S7:**
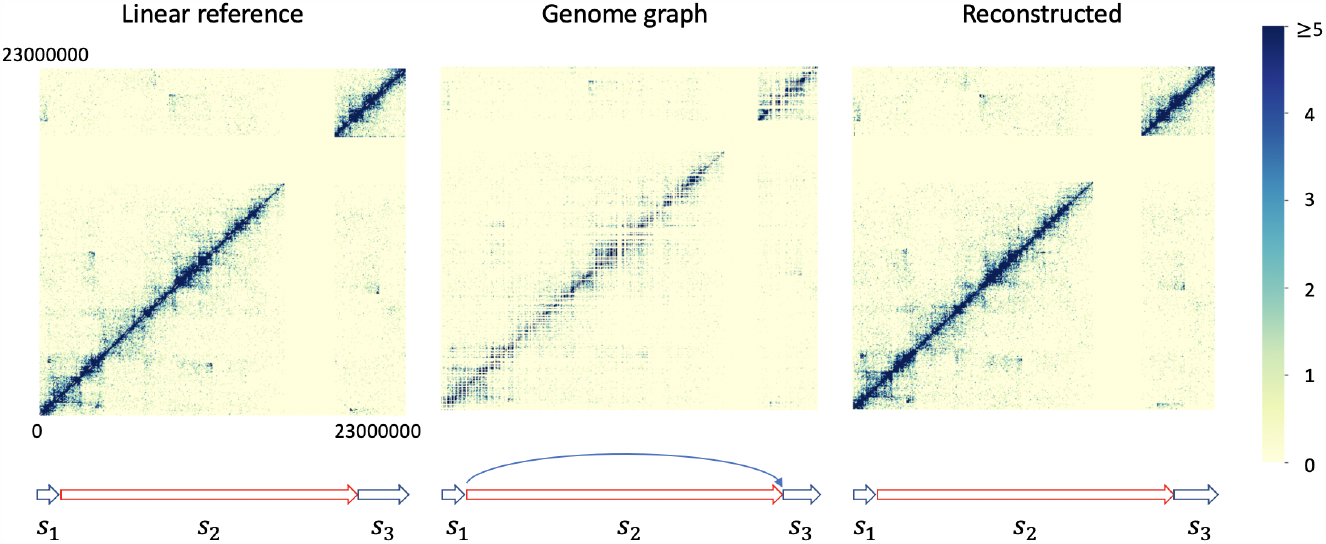
The region between 0bp and 23, 000, 000bp in chromosome 18. The genome graph shows a large deletion (*s*_2_) with an approximate length of 20, 000, 000bp. The empty stripe is the centromere region.

**Table S4:**
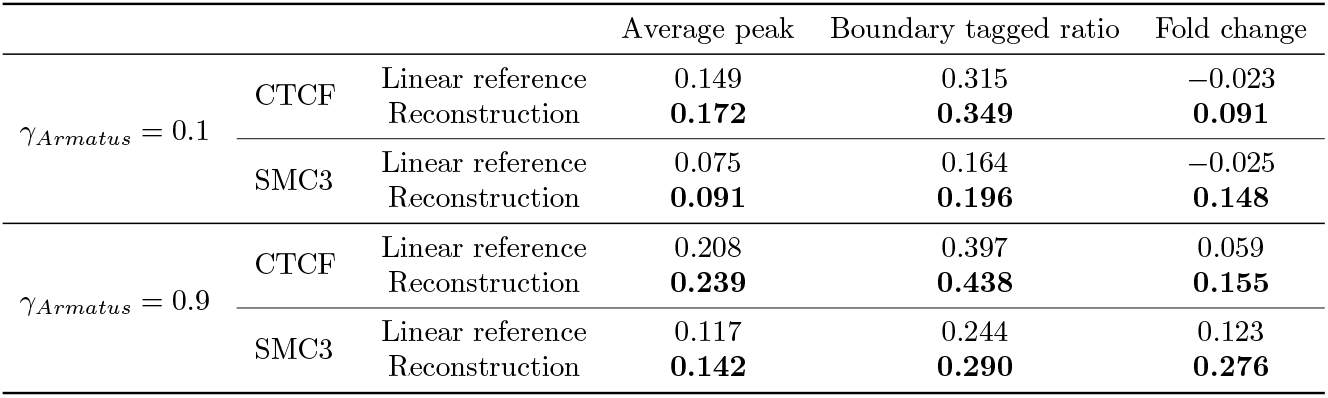
The comparisons of three metrics reflecting CTCF or SMC3 enrichments near TAD boundaries across different genomes. *γ*_*Armatus*_ denotes the hyper-parameter *γ* of the TAD caller Armatus. Hi-C sample: SRR1659693.

**Figure S8:**
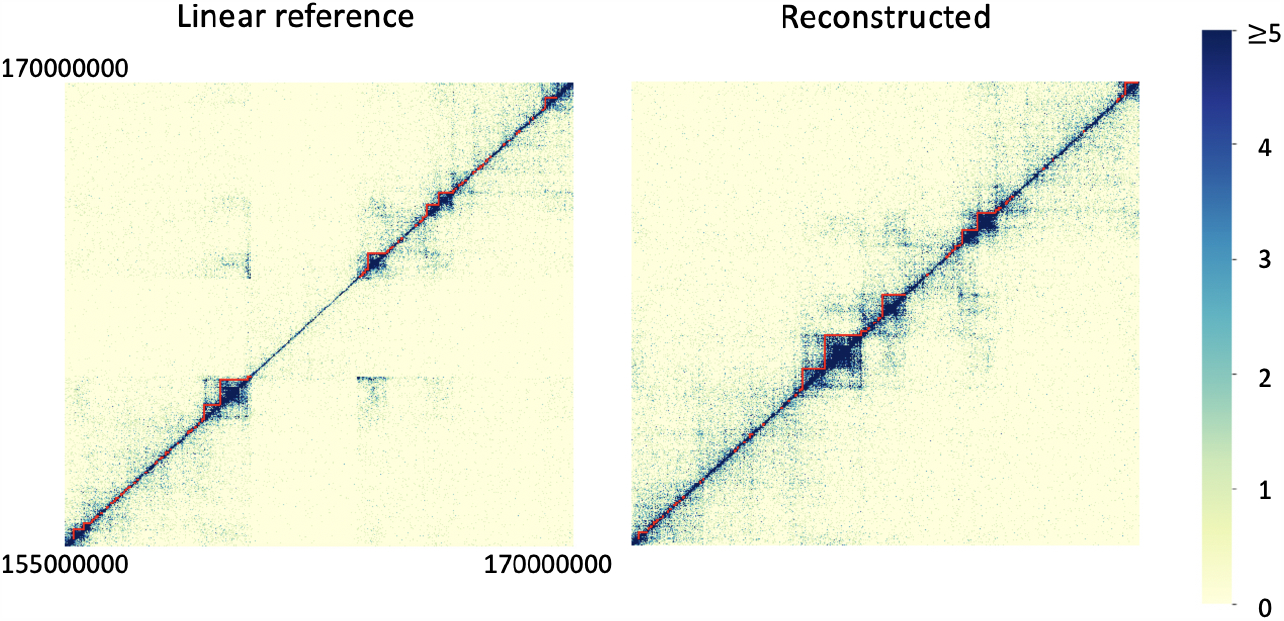
TADs in the same region as Figure 2. TADs are called by Armatus with hyper-parameter *γ*_*Armatus*_ = 0.5.

**Figure S9:**
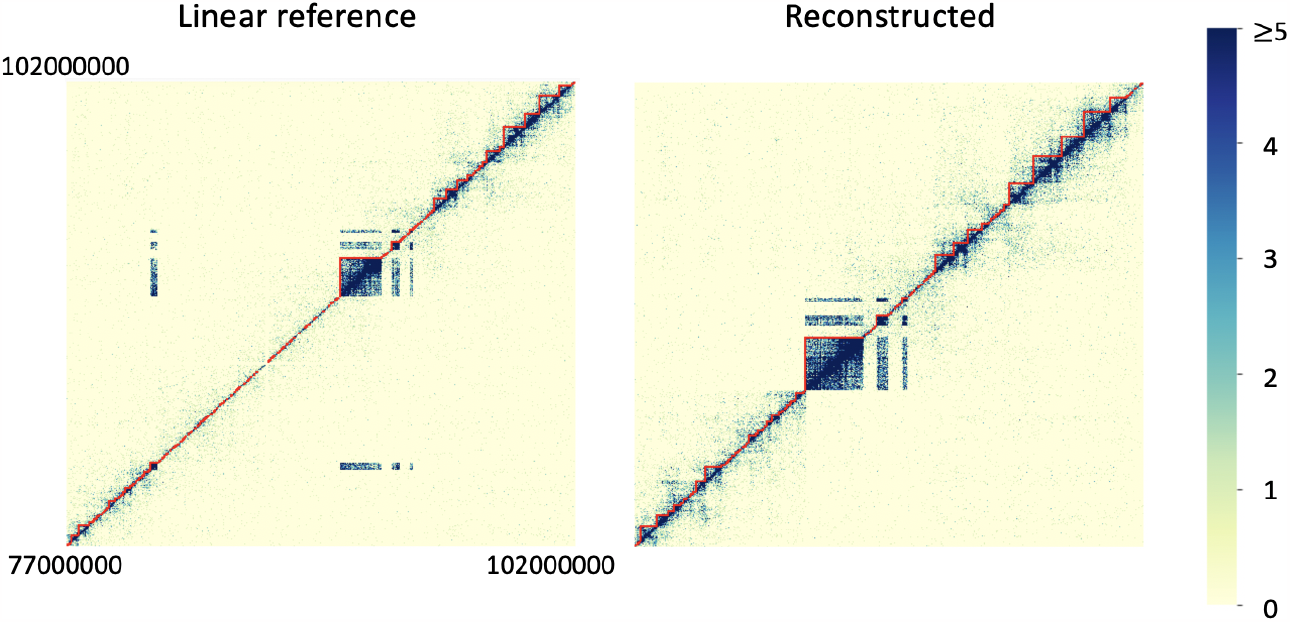
TADs in the same region as Figure S5. TADs are called by Armatus with hyper-parameter *γ*_*Armatus*_ = 0.5.

**Figure S10:**
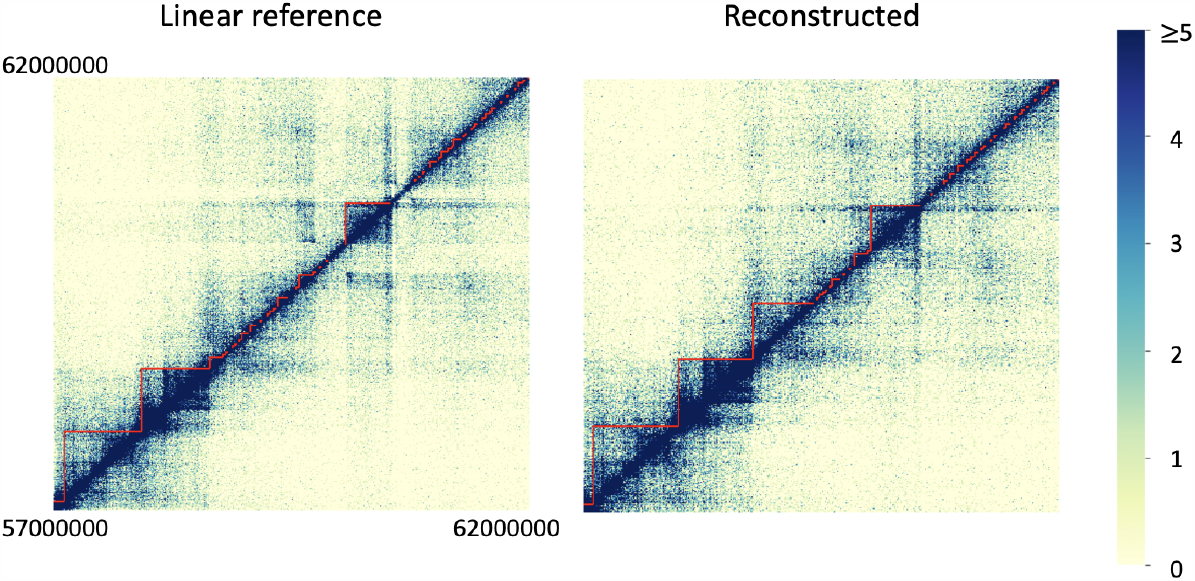
TADs in the same region as Figure 3. TADs are called by Armatus with hyper-parameter *γ*_*Armatus*_ = 0.5.

**Table S5:** The comparisons of three metrics reflecting CTCF or SMC3 enrichments near TAD boundaries across different genomes. *γ*_*Armatus*_ denotes the hyper-parameter *γ* of the TAD caller Armatus. Hi-C sample: SRR1659694.

|  |  |  | Average peak | Boundary tagged ratio | Fold change |
| --- | --- | --- | --- | --- | --- |
| $\gamma_{\text{Armatus}} = 0.1$ | CTCF | Linear reference | 0.152 | 0.320 | -0.008 |
|  |  | Reconstruction | **0.177** | **0.356** | **0.109** |
|  | SMC3 | Linear reference | 0.076 | 0.168 | 0.005 |
|  |  | Reconstruction | **0.096** | **0.203** | **0.197** |
| $\gamma_{\text{Armatus}} = 0.5$ | CTCF | Linear reference | 0.175 | 0.350 | 0.037 |
|  |  | Reconstruction | **0.213** | **0.403** | **0.181** |
|  | SMC3 | Linear reference | 0.093 | 0.198 | 0.076 |
|  |  | Reconstruction | **0.121** | **0.246** | **0.292** |
| $\gamma_{\text{Armatus}} = 0.9$ | CTCF | Linear reference | 0.212 | 0.403 | 0.084 |
|  |  | Reconstruction | **0.254** | **0.457** | **0.201** |
|  | SMC3 | Linear reference | 0.120 | 0.249 | 0.149 |
|  |  | Reconstruction | **0.152** | **0.306** | **0.338** |

**Figure S11:**
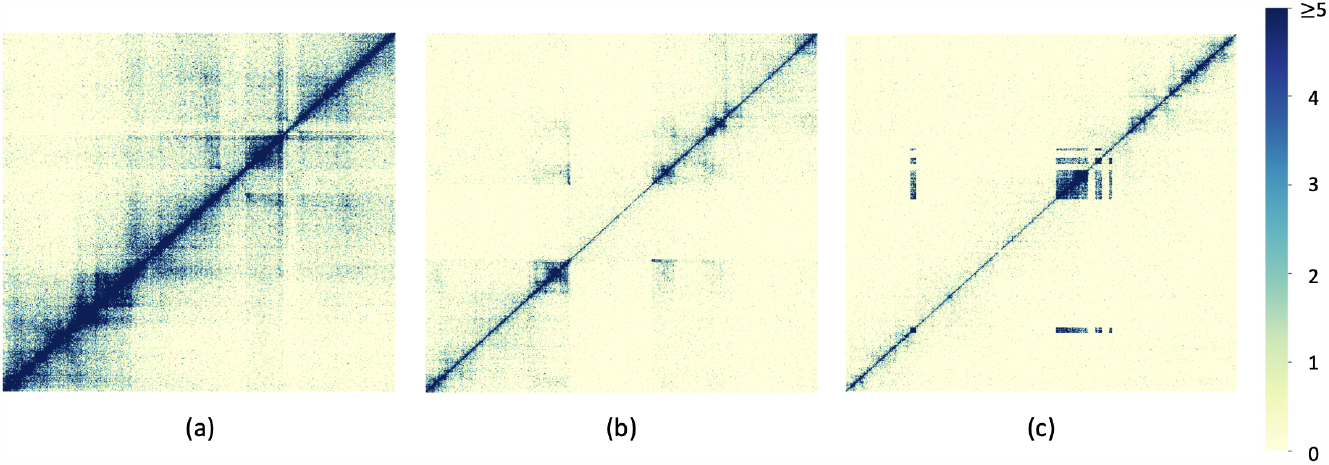
Three regions of contact matrices generated by the longest *M* -weighted path algorithm, corresponding to regions shown in Figure 3, Figure 2, and Figure S5.

**Table S6:** The values of three metrics reflecting CTCF or SMC3 enrichments near TAD boundaries generated directly from dynamic programming.

|  |  | Average peak | Boundary tagged ratio | Fold change |
| --- | --- | --- | --- | --- |
| TADs from DP | CTCF | 0.282 | 0.522 | 0.070 |
|  | SMC3 | 0.164 | 0.348 | 0.113 |

**Table S7:** Mapping statistics of Hi-C reads being mapped onto different references, computed by `vg stats -a`. Shortest path: reads mapped onto the linear genome inferred by shortest path algorithm; Longest *M* -weighted path: reads mapped onto the linear genome inferred by longest *M* -weighted path algorithm.

| SRR1658693 |  |  |
| --- | --- | --- |
| | Shortest path | Longest $M$-weighted path |
| Mapped reads | 873,813,004 | 878,342,257 |
| Perfectly mapped reads | 286,269,146 | 289,729,014 |

**Table S8:** Mapping statistics of Hi-C reads being mapped onto different references, computed by `vg stats -a`. Shortest path: reads mapped onto the linear genome inferred by shortest path algorithm; Longest *M* -weighted path: reads mapped onto the linear genome inferred by longest *M* -weighted path algorithm.

| SRR1658694 |  |  |
| --- | --- | --- |
| | Shortest path | Longest $M$-weighted path |
| Mapped reads | 1,161,272,502 | 1,166,868,005 |
| Perfectly mapped reads | 598,340,773 | 605,592,165 |

**Table S9:**
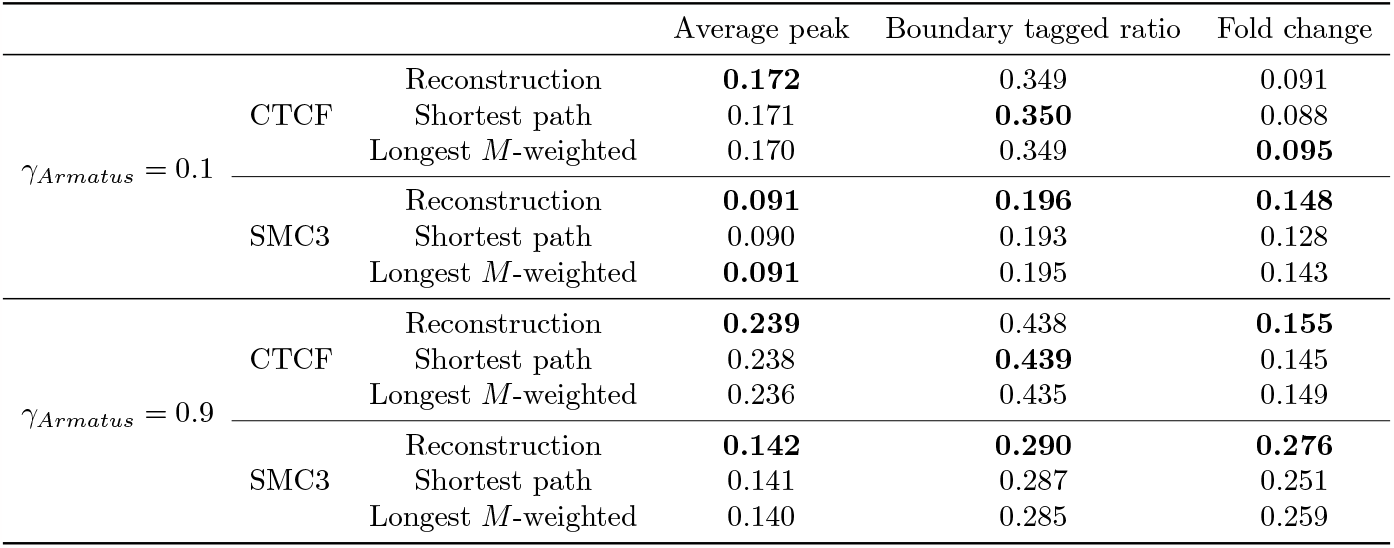
The comparisons of three metrics reflecting CTCF or SMC3 enrichments near TAD boundaries across different genomes. Reconstruction: the genome inferred by our algorithm. Shortest path: the genome inferred by shortest path algorithm; Longest *M* -weighted: the genome inferred by longest *M* - weighted path algorithm. *γ*_*Armatus*_ denotes the hyper-parameter *γ* of the TAD caller Armatus. Hi-C sample: SRR1659693.

## References

Peter Fraser and Wendy Bickmore. Nuclear organization of the genome and the potential for gene regulation. Nature, 447(7143):413–417, 2007.

Sarah Rennie, Maria Dalby, Lucas van Duin, and Robin Andersson. Transcriptional decomposition re-veals active chromatin architectures and cell specific regulatory interactions. Nature Communications, 9(1):487, 2018.

Shiv IS Grewal and Danesh Moazed. Heterochromatin and epigenetic control of gene expression. Science, 301(5634):798–802, 2003.

Benjamin D Pope, Tyrone Ryba, Vishnu Dileep, Feng Yue, Weisheng Wu, Olgert Denas, Daniel L Vera, Yanli Wang, R Scott Hansen, Theresa K Canfield, et al. Topologically associating domains are stable units of replication-timing regulation. Nature, 515(7527):402–405, 2014.

Erez Lieberman-Aiden, Nynke L Van Berkum, Louise Williams, Maxim Imakaev, Tobias Ragoczy, Agnes Telling, Ido Amit, Bryan R Lajoie, Peter J Sabo, Michael O Dorschner, et al. Comprehensive mapping of long-range interactions reveals folding principles of the human genome. Science, 326(5950):289–293, 2009.

Job Dekker, Karsten Rippe, Martijn Dekker, and Nancy Kleckner. Capturing chromosome conformation. Science, 295(5558):1306–1311, 2002.

Elphège P Nora, Bryan R Lajoie, Edda G Schulz, Luca Giorgetti, Ikuhiro Okamoto, Nicolas Servant, Tristan Piolot, Nynke L Van Berkum, Johannes Meisig, John Sedat, et al. Spatial partitioning of the regulatory landscape of the X-inactivation centre. Nature, 485(7398):381–385, 2012.

Jesse R Dixon, Siddarth Selvaraj, Feng Yue, Audrey Kim, Yan Li, Yin Shen, Ming Hu, Jun S Liu, and Bing Ren. Topological domains in mammalian genomes identified by analysis of chromatin interactions. Nature, 485(7398):376–380, 2012.

Wouter De Laat and Denis Duboule. Topology of mammalian developmental enhancers and their regulatory landscapes. Nature, 502(7472):499–506, 2013.

Darya Filippova, Rob Patro, Geet Duggal, and Carl Kingsford. Identification of alternative topological domains in chromatin. Algorithms for Molecular Biology, 9:1–11, 2014.

Angsheng Li, Xianchen Yin, Bingxiang Xu, Danyang Wang, Jimin Han, Yi Wei, Yun Deng, Ying Xiong, and Zhihua Zhang. Decoding topologically associating domains with ultra-low resolution Hi-C data by graph structural entropy. Nature Communications, 9(1):3265, 2018.

Abbas Roayaei Ardakany, Halil Tuvan Gezer, Stefano Lonardi, and Ferhat Ay. Mustache: multi-scale detection of chromatin loops from Hi-C and Micro-C maps using scale-space representation. Genome Biology, 21:1–17, 2020.

Su Wang, Soohyun Lee, Chong Chu, Dhawal Jain, Peter Kerpedjiev, Geoffrey M Nelson, Jennifer M Walsh, Burak H Alver, and Peter J Park. HiNT: a computational method for detecting copy number variations and translocations from Hi-C data. Genome Biology, 21:1–15, 2020.

Robert Schöpflin, Uirá Souto Melo, Hossein Moeinzadeh, David Heller, Verena Laupert, Jakob Hertzberg, Manuel Holtgrewe, Nico Alavi, Marius-Konstantin Klever, Julius Jungnitsch, et al. Integration of Hi-C with short and long-read genome sequencing reveals the structure of germline rearranged genomes. Nature Communications, 13(1):6470, 2022.

Ahmed Ibrahim Samir Khalil, Siti Rawaidah Binte Mohammad Muzaki, Anupam Chattopadhyay, and Amartya Sanyal. Identification and utilization of copy number information for correcting Hi-C contact map of cancer cell lines. BMC Bioinformatics, 21(1):1–25, 2020.

Xiaotao Wang, Jie Xu, Baozhen Zhang, Ye Hou, Fan Song, Huijue Lyu, and Feng Yue. Genome-wide detection of enhancer-hijacking events from chromatin interaction data in rearranged genomes. Nature Methods, 18(6):661–668, 2021.

Ying Gong, Yefang Li, Xuexue Liu, Yuehui Ma, and Lin Jiang. A review of the pangenome: how it affects our understanding of genomic variation, selection and breeding in domestic animals? Journal of Animal Science and Biotechnology, 14(1):1–19, 2023.

Ting Wang, Lucinda Antonacci-Fulton, Kerstin Howe, Heather A Lawson, Julian K Lucas, Adam M Phillippy, Alice B Popejoy, Mobin Asri, Caryn Carson, Mark JP Chaisson, et al. The Human Pangenome Project: a global resource to map genomic diversity. Nature, 604(7906):437–446, 2022.

Adam Ameur. Goodbye reference, hello genome graphs. Nature Biotechnology, 37(8):866–868, 2019.

Erik Garrison, Jouni Sirén, Adam M Novak, Glenn Hickey, Jordan M Eizenga, Eric T Dawson, William Jones, Shilpa Garg, Charles Markello, Michael F Lin, et al. Variation graph toolkit improves read mapping by representing genetic variation in the reference. Nature Biotechnology, 36(9):875–879, 2018.

Yutong Qiu and Carl Kingsford. Constructing small genome graphs via string compression. Bioinformatics, 37(Supplement 1):i205–i213, 2021.

Glenn Hickey, Jean Monlong, Jana Ebler, Adam M Novak, Jordan M Eizenga, Yan Gao, Tobias Marschall, Heng Li, and Benedict Paten. Pangenome graph construction from genome alignments with Minigraph-Cactus. Nature Biotechnology, pages 1–11, 2023.

Erik Garrison, Andrea Guarracino, Simon Heumos, Flavia Villani, Zhigui Bao, Lorenzo Tattini, Jörg Hagmann, Sebastian Vorbrugg, Santiago Marco-Sola, Christian Kubica, et al. Building pangenome graphs. bioRxiv, pages 2023–04, 2023.

Prashant Pandey, Yinjie Gao, and Carl Kingsford. VariantStore: an index for large-scale genomic variant search. Genome Biology, 22(1):1–25, 2021.

Goran Rakocevic, Vladimir Semenyuk, Wan-Ping Lee, James Spencer, John Browning, Ivan J Johnson, Vladan Arsenijevic, Jelena Nadj, Kaushik Ghose, Maria C Suciu, et al. Fast and accurate genomic analyses using genome graphs. Nature Genetics, 51(2):354–362, 2019.

Daehwan Kim, Joseph M Paggi, Chanhee Park, Christopher Bennett, and Steven L Salzberg. Graph-based genome alignment and genotyping with HISAT2 and HISAT-genotype. Nature Biotechnology, 37(8):907–915, 2019.

Mikko Rautiainen and Tobias Marschall. GraphAligner: rapid and versatile sequence-to-graph alignment. Genome Biology, 21(1):253, 2020.

Jouni Sirén, Jean Monlong, Xian Chang, Adam M Novak, Jordan M Eizenga, Charles Markello, Jonas A Sibbesen, Glenn Hickey, Pi-Chuan Chang, Andrew Carroll, et al. Pangenomics enables genotyping of known structural variants in 5202 diverse genomes. Science, 374(6574):abg8871, 2021.

Chen-Shan Chin, Sairam Behera, Asif Khalak, Fritz J Sedlazeck, Peter H Sudmant, Justin Wagner, and Justin M Zook. Multiscale analysis of pangenomes enables improved representation of genomic diversity for repetitive and clinically relevant genes. Nature Methods, pages 1–9, 2023.

Kevin Hadi, Xiaotong Yao, Julie M Behr, Aditya Deshpande, Charalampos Xanthopoulakis, Huasong Tian, Sarah Kudman, Joel Rosiene, Madison Darmofal, Joseph DeRose, et al. Distinct classes of complex structural variation uncovered across thousands of cancer genome graphs. Cell, 183(1):197–210, 2020.

Hannes P Eggertsson, Snaedis Kristmundsdottir, Doruk Beyter, Hakon Jonsson, Astros Skuladottir, Marteinn T Hardarson, Daniel F Gudbjartsson, Kari Stefansson, Bjarni V Halldorsson, and Pall Melsted. GraphTyper2 enables population-scale genotyping of structural variation using pangenome graphs. Nature Communications, 10(1):5402, 2019.

Sai Chen, Peter Krusche, Egor Dolzhenko, Rachel M Sherman, Roman Petrovski, Felix Schlesinger, Melanie Kirsche, David R Bentley, Michael C Schatz, Fritz J Sedlazeck, et al. Paragraph: a graph-based structural variant genotyper for short-read sequence data. Genome Biology, 20:1–13, 2019.

Wen-Wei Liao, Mobin Asri, Jana Ebler, Daniel Doerr, Marina Haukness, Glenn Hickey, Shuangjia Lu, Julian K Lucas, Jean Monlong, Haley J Abel, et al. A draft human pangenome reference. Nature, 617 (7960):312–324, 2023.

Xiao-Tao Wang, Wang Cui, and Cheng Peng. HiTAD: detecting the structural and functional hierarchies of topologically associating domains from chromatin interactions. Nucleic Acids Research, 45(19):e163–e163, 2017.

Ferhat Ay, Timothy L Bailey, and William Stafford Noble. Statistical confidence estimation for Hi-C data reveals regulatory chromatin contacts. Genome Research, 24(6):999–1011, 2014.

Nicolas Servant, Nelle Varoquaux, Bryan R Lajoie, Eric Viara, Chong-Jian Chen, Jean-Philippe Vert, Edith Heard, Job Dekker, and Emmanuel Barillot. HiC-Pro: an optimized and flexible pipeline for Hi-C data processing. Genome Biology, 16(1):1–11, 2015.

Harold N. Gabow, Shachindra N Maheshwari, and Leon J. Osterweil. On two problems in the generation of program test paths. IEEE Transactions on Software Engineering, (3):227–231, 1976.

Petr Kolman and Ond řej Pangrác. On the complexity of paths avoiding forbidden pairs. Discrete Applied Mathematics, 157(13):2871–2876, 2009.

Caxton C Foster. A generalization of AVL trees. Communications of the ACM, 16(8):513–517, 1973.

Boyan Bonev and Giacomo Cavalli. Organization and function of the 3D genome. Nature Reviews Genetics, 17(11):661–678, 2016.

Bo Zhou, Steve S Ho, Stephanie U Greer, Xiaowei Zhu, John M Bell, Joseph G Arthur, Noah Spies, Xianglong Zhang, Seunggyu Byeon, Reenal Pattni, et al. Comprehensive, integrated, and phased whole-genome analysis of the primary ENCODE cell line K562. Genome Research, 29(3):472–484, 2019.

Suhas SP Rao, Miriam H Huntley, Neva C Durand, Elena K Stamenova, Ivan D Bochkov, James T Robinson, Adrian L Sanborn, Ido Machol, Arina D Omer, Eric S Lander, et al. A 3d map of the human genome at kilobase resolution reveals principles of chromatin looping. Cell, 159(7):1665–1680, 2014.

Ivar Grytten, Knut D Rand, Alexander J Nederbragt, Geir O Storvik, Ingrid K Glad, and Geir K Sandve. Graph Peak Caller: Calling ChIP-seq peaks on graph-based reference genomes. PLoS computational biology, 15(2):e1006731, 2019.

Jouni Sirén. Indexing variation graphs. In 2017 Proceedings of the ninteenth workshop on algorithm engineering and experiments (ALENEX), pages 13–27. SIAM, 2017.

Taku Onodera, Kunihiko Sadakane, and Tetsuo Shibuya. Detecting superbubbles in assembly graphs. In International workshop on algorithms in bioinformatics, pages 338–348. Springer, 2013.

Bryan R Lajoie, Job Dekker, and Noam Kaplan. The Hitchhiker’s guide to Hi-C analysis: practical guidelines. Methods, 72:65–75, 2015.

Elarbi Choukhmane and John Franco. An approximation algorithm for the maximum independent set problem in cubic planar graphs. Networks, 16(4):349–356, 1986.

Helmuth Späth. One dimensional spline interpolation algorithms. CRC press, 1995.

Marie Zufferey, Daniele Tavernari, Elisa Oricchio, and Giovanni Ciriello. Comparison of computational methods for the identification of topologically associating domains. Genome Biology, 19(1):1–18, 2018.

Kun Liu, Hong-Dong Li, Yaohang Li, Jun Wang, and Jianxin Wang. A comparison of topologically associating domain callers based on Hi-C data. IEEE/ACM Transactions on Computational Biology and Bioinformatics, 20(1):15–29, 2022.

Emre Sefer. A comparison of topologically associating domain callers over mammals at high resolution. BMC Bioinformatics, 23(1):127, 2022.

